# Variation in a microbial mutualist has transcriptional and phenotypic consequences for host-parasite interactions

**DOI:** 10.1101/2024.07.15.603596

**Authors:** Addison D. Buxton-Martin, Emile Gluck-Thaler, Eunnuri Yi, John R. Stinchcombe, Corlett W. Wood

## Abstract

Strains of microbial symbionts often vary in their effect on their host. However, little is known about how the genetic variation in microbial symbiont populations impacts host interactions with other co-colonizing microbes. We investigated how different strains of nitrogen-fixing rhizobial bacteria affect their host plant’s response to parasitic root-knot nematodes in the legume *Medicago truncatula*. Using dual RNAseq of the root organs harboring rhizobia or nematodes, we identified genes from plant host, rhizobia, and nematode whose expression differed between parasite-infected and-uninfected plants, and between plants inoculated with different rhizobial strains. At the site of host-parasite interactions (in nematode galls), hundreds of host genes and few nematode genes differed in expression between host plants inoculated with different rhizobia strains. At the site of host-mutualist interactions (in rhizobia nodules), hundreds of host genes and few rhizobial genes responded to parasite infection. The vast majority of parasite-induced changes in host gene expression depended on the resident rhizobia strain. We additionally observed differences in parasite load and in some root architecture traits between plants inoculated with different rhizobia strains, showing that genetic variation in a mutualistic symbiont impacted parasite colonization. The transcriptomic and phenotypic differences we observed suggest that microbial indirect genetic effects play an underappreciated role in their host’s interactions with other co-colonizing microorganisms.

## Introduction

An individual plant hosts thousands of microorganisms, ranging from bacteria and viruses to eukaryotes like fungi and protozoa. These microbial species are composed of strains that vary – often dramatically – in their effects on their host (Heath, 2010; Heath and Stinchcombe, 2014; Lange *et al*., 2023; Lively and Dybdahl, 2000; Morand *et al*., 1997; Osborne *et al*., 2009; Parker *et al*., 2017). Here we investigate how strain-level genetic variation in microbial symbiont populations cascades to affect a plant’s interactions with other microbes.

Co-colonizing symbionts – parasites, commensals, or mutualists – often compete with or facilitate one another (Coyte *et al*., 2015; Stein *et al*., 2013) directly or indirectly via their effects on their shared host (Foster *et al*., 2017; Pedersen and Fenton, 2007). This framework is implicitly or explicitly employed by researchers studying interactions between hosts and microbial communities in a variety of disciplines: in plant pathology, as the priming of host defenses (Ahmad *et al*., 2010; Benjamin *et al*., 2022; Guo and Ge, 2017; Zamioudis and Pieterse, 2012); in parasitology within the context of co-infections or mixed infections (Alizon *et al*., 2013; Devi *et al*., 2021; Venter *et al*., 2022); and in community and microbial ecology, as priority effects, indirect effects, or higher-order interactions (Debray *et al*., 2022; Miller and Travis, 1996; Strauss, 1991). Across these disparate subfields, consensus has emerged that interactions between co-colonizing microbes of different species have far-reaching ramifications for the assembly and function of host-associated communities (Alizon *et al*., 2013; Banerjee *et al*., 2018; Dastogeer *et al*., 2020; Shetty *et al*., 2017; terHorst *et al*., 2018). Co-colonizing microbes of different species can influence the relationship their co-colonizers have with their joint host at all stages of their respective lifecycles and in different ways: from impacting each other’s colonization success (Telfer *et al*., 2010), transmission dynamics (Bousema *et al*., 2008; Pollitt *et al*., 2015; Schneider *et al*., 2018), or the benefits they confer and costs they inflict on their shared host (Barrett *et al*., 2015; Larimer *et al*., 2014; Magnoli and Bever, 2023; Weldon *et al*., 2020).

However, within symbiont species there is extensive genetic and functional variation. For example, pathogen genotypes differ dramatically in virulence and transmissibility (Lively and Dybdahl, 2000; Morand *et al*., 1997), strains of bacterial mutualists vary in the costs and benefits they bring to their hosts (Heath, 2010; Heath and Stinchcombe, 2014; Heath and Tiffin, 2007; Calvert *et al*., 2023), and genotypes of defensive mutualists differ in their protective effect (Lange *et al*., 2023; Osborne *et al*., 2009; Parker *et al*., 2017). These examples share a unifying feature: the genes in one species (the microbe) impact trait expression in another (the host). Such “interspecific indirect genetic effects” (De Lisle *et al*., 2022; Shuster *et al*., 2006) are thought to be pervasive in traits mediating symbiotic interactions (Fronk and Sachs, 2022) and alter the tempo and mode of trait evolution, because the genetic basis of trait expression and evolution in one species (the host) resides in another species (the microbe) (De Lisle *et al*., 2022; O’Brien *et al*., 2021b; Queller, 2014; Shuster *et al*., 2006).

The extensive variation of symbiosis traits within microbial species raises a crucial question: do different microbial strains of the same species vary in their effect on the host’s interaction with other microbes? Put another way: can genetic variation for traits mediating species interactions reside in species not directly involved in the interaction itself? The answer to this question has broad implications for the assembly, function, and evolution of host-associated communities. If strain-level (i.e., intraspecific) variation in a microbial symbiont influences its host’s interaction with other symbionts, then strain-level changes in microbial symbiont populations such as changes in allele frequencies or strain abundance could have cascading impacts on the structure or function of the entire host-associated community. The few studies that have addressed this possibility have found that microbial genotypes do vary in their effect on the broader host-associated community, especially in defensive mutualisms (Lange *et al*., 2023; Osborne *et al*., 2009; Parker *et al*., 2017). However, when two microbes inhabit the same niche within the host (e.g., the animal gut) or colonize systemically, disentangling the host’s responses to two co-colonizing symbionts and the indirect effects between symbionts can be challenging.

Plant galls provide a unique means of circumventing these challenges and investigating how genetic variation in one microbial symbiont species impacts its host’s interactions with other co-colonizing symbionts. Galls are specialized structures that are made of plant tissue but are induced and colonized by another organism such as an insect or microbe. Many plant parasites, pathogens, and mutualists form galls (Harris and Pitzschke, 2020). Common examples include legume root nodules induced by nitrogen-fixing rhizobia, goldenrod galls induced by the goldenrod gall fly, and root knot galls induced by root-knot nematodes (Harris and Pitzschke, 2020). These galls are the locus of interaction between the gall inducers and their host plant (Fronk and Sachs, 2022). Because each gall is specific to a single pairwise species interaction, galls provide a unique opportunity to sample the symbiotic interface between the host and single symbiont even in hosts interacting with many other symbionts. The abundance and discrete nature of these structures is also helpful in quantifying and phenotyping aspects of host-symbiont interactions such as parasite or mutualist colonization rates or host susceptibility.

Transcriptomic methods when applied to plant galls are particularly useful in elucidating the otherwise difficult to observe interaction involving hosts and microbes, as they provide genome-wide proxies of the underlying biology mediating the interaction between host and symbiont. In particular, dual RNAseq, where RNA from two symbiotic organisms are sequenced in the same sample, has become a useful tool to investigate in vivo interactions between a host and symbiont (Westermann *et al*., 2017; Westermann and Vogel, 2021; Wolf *et al*., 2018). Thus comparing the gene expression profiles of galls can be used to examine changes in the interactions between host plants and their microbial gall-forming symbionts under different conditions. What is more, interactions between plants and symbiotic microbes may drive systemic reactions outside of just the organ that mediates the host-symbiont interaction (Pieterse *et al*., 2014; Mauch-Mani *et al*., 2017) and plant-microbe interactions can often be regulated by signals produced in parts of the host plant not directly involved in the interaction (Li *et al*., 2022). Comparing gene expression profiles of galls to that of other organs can thus be used to assess the systemic or local nature of an interaction between host and symbiont (Oates *et al.,* 2021).

We leveraged these unique features of root-knot nematode galls and rhizobia nodules as well as dual RNAseq to investigate how strain-level variation in a microbial mutualist influences its host plant’s transcriptomic response to parasite infection. We inoculated a single genotype of the model legume *Medicago truncatula* with one of two strains of mutualistic nitrogen-fixing rhizobia bacteria *Ensifer meliloti* and infected half of these plants with the parasitic root-knot nematode *Meloidogyne hapla*. We then profiled plant, mutualist, and parasite gene expression in rhizobia nodules and nematode galls (Figure **1**) and quantified host fitness, root architecture, and parasite and mutualist colonization rates. We asked: (1) how do mutualistic rhizobia strains influence host traits, parasite colonization rates, and gene expression in nematode galls?; (2) how do parasitic nematodes affect host traits, mutualist colonization rates, and gene expression in rhizobial nodules?; and (3) is the effect of parasite infection on host traits, mutualist colonization rates, and nodule gene expression influenced by the resident rhizobia strain?

**Figure 1:**
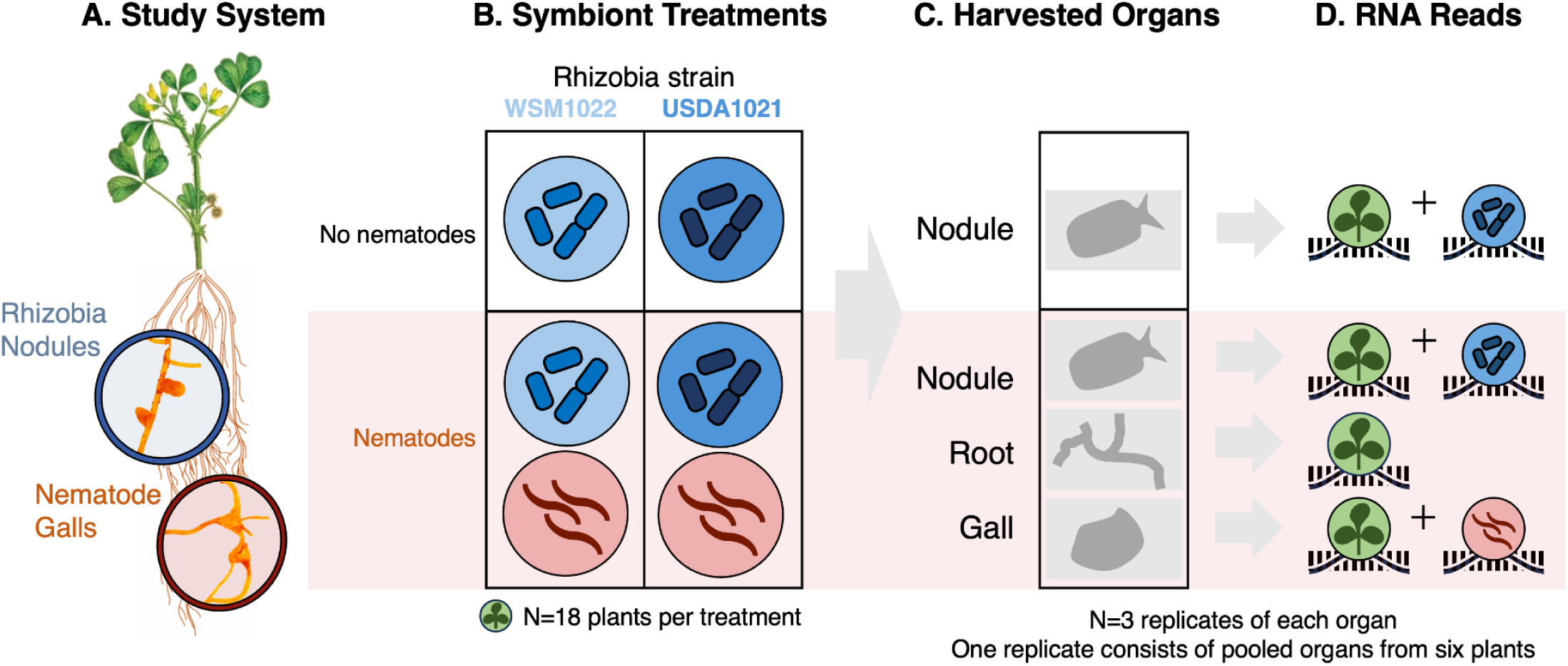
Conceptual diagram of the experimental design, treatments, and samples used in this experiment. A) The host plant, *Medicago truncatula*, with rhizobia nodules and nematode galls in the host plant root system as magnified insets. B) Each of the four treatments of symbionts consisted of three biological replicates with six plants constituting each biological replicate, totaling 18 plants per treatment. C) Rhizobia nodules, roots, and nematode galls were harvested from hosts as indicated. Organs from the six plants constituting one biological replicate were pooled. D) The recoverable RNA reads from each organ.

On a phenotypic level, we found that nematode infection status impacted root architecture and rhizobia colonization rates, while rhizobia strain impacted parasite colonization rate. Neither nematode infection status, nematode infection severity, or rhizobia strain impacted a host fitness proxy, whereas rhizobia colonization rates increased host fitness. On a transcriptomic level, we found that strain-level variation in the rhizobial mutualist significantly alters the host plant’s transcriptomic response to parasite infection. Plants inoculated with different rhizobial strains had differential gene expression in nematode galls, the site of parasite infection.

Furthermore, parasite infection altered the expression of hundreds of genes in rhizobia nodules, and this effect depended strongly on the resident rhizobia strain. Our results show that intraspecific genetic variation in a keystone microbial mutualist has far reaching consequences on host-parasite interactions with mutualist strain having transcriptional consequences in mature host-parasite structures and consequences for parasite colonization rates. What is more, parasite infection carried consequences for host-mutualist interactions with transcriptional consequences in mature host-mutualist structures and altered mutualist colonization rates.

## Materials and Methods

### Replication Statement

| Scale of inference | The scale at which the factor of interest is applied | Number of replicates at the appropriate scale |
| --- | --- | --- |
| Population of plants (transcriptomic response) | Manipulated parasitic nematode infection and rhizobia strain inoculation on individual plants; pooled organs (gall, nodule, or root) from 6 individual plants into a single biological replicate | 3 replicates of each organ per treatment (nematode infection and rhizobia strain combination) |
| Population of plants (phenotypic response) | Manipulated parasitic nematode infection and rhizobia strain inoculation on individual plants | 19 plants per combination of nematode treatment and rhizobia strain |

### Study system

*Medicago truncatula* Gaertn. is an annual legume native to the Mediterranean (Cook, 1999). Upon the transfer of compatible chemical signals, *M. truncatula* plants form associations with nitrogen-fixing rhizobial bacteria in the soil, such as *Ensifer meliloti*. Rhizobia are found throughout soil and most legumes are colonized by rhizobia in the wild. Rhizobia are housed in nodules that develop from the roots of their host plant. Nodules provide the requisite low-oxygen environment and resources for the rhizobia to differentiate into bacteroids within intracellular compartments called symbiosomes. Within these symbiosomes, rhizobia fix atmospheric nitrogen into biologically available forms of nitrogen and provide some to their host plant and in exchange hosts provide rhizobia with carbon-containing compounds derived from photosynthate. The *Medicago*-*Ensifer* system has become a well characterized model of resource-exchange mutualisms (Long, 1989).

The pathogenic root-knot nematode *Meloidogyne hapla* also associates with *M. truncatula* roots. Female *M. hapla* burrow through the intercellular spaces of host roots in search of a suitable infection site. There *M. hapla* uses its stylet mouthpiece to inject effector proteins into parenchymal host cells and root tissues that suppress immune responses and induce cell differentiation into giant cells from which the pathogenic nematode siphons nutrients. Nematode galls form around this site of infection (Jones and Goto, 2011).

We chose the in-bred line of *M. truncatula* accession HM145 from the French National Institute for Agriculture Research (INRAE) germplasm library for its high nodulation rate and nematode susceptibility (Wood *et al*., 2018). We obtained inoculum containing the northern root-knot nematode *M. hapla* line LM originally collected in France and supplied by the Department of Entomology and Plant Pathology at North Carolina State University. We used *E. meliloti* strains USDA1021 and WSM1022 from lab stocks (JRS). WSM1022 is known as a high-efficiency nitrogen fixer in contrast to other strains (Terpolilli *et al*., 2013).

### Sample Preparation

We grew *M. truncatula* HM145 in Cone-tainers and maintained them in growth chambers at [university removed for anonymity] following the instructions of Per Garcia *et al*. (2006).

Plants were arranged in a randomized complete block design of rhizobia strain (USDA1021 or WSM1022) and nematode status (infected or uninfected) and plants were inoculated with the appropriate nematode and rhizobia treatment on day 14 after planting. Each treatment had three replicates, which were each composed of six individual plants. See Methods S1 and Table S3 for greater detail regarding sample preparation. We did not include a nematode-only treatment because a nematode-only treatment would be ecologically unrealistic: rhizobia colonization of legumes is pervasive in natural settings, meaning that nearly all *M. truncatula* that are infected with *M. hapla* are also colonized with *E. meliloti*. Furthermore *M. truncatula* that are not colonized by *E. meliloti* suffer from significant nutrient deficit, making it nearly impossible to disentangle the effects of nematode infection alone from nutrient deficiency.

### Organ sample harvesting

Organ samples were collected from all plants 52-53 days after planting (see Methods S1 for rationale regarding this timeframe). All six plants that constituted a replicate were harvested on the same day. Nodule samples were collected from all plants and root and nematode gall samples were collected from plants inoculated with both rhizobia and nematodes (Figure **1**). To harvest, the root system was washed in water, then mature organs were manually excised using a pair of sterilized scissors and immediately flash frozen in liquid nitrogen. Samples were taken from the top ⅓ of the root system so as to collect nematode galls and nodules at a consistent level of maturity. All samples were stored at-80 °C until RNA extraction which was performed within 30 days of harvesting.

### Dual RNA sequencing of nodules and nematode galls and RNA sequencing of roots

To isolate species interactions in co-colonized hosts, we extracted RNA from nodules and nematode galls for Dual RNAseq. To assess the localized or systemic nature of host changes in gene expression, we also extracted RNA from samples of roots from co-colonized hosts. We used a Trizol and QIAGEN RNeasy Kit based RNA extract and a Nanodrop ND-1000 (Thermo Fisher Scientific) to confirm RNA quality. RNA extracts were shipped to Genewiz Azenta Life Sciences (South Plainfield, NJ) for sequencing. cDNA preparation was performed by Genewiz Azenta Life Sciences using NEBNext Ultra II RNA Library Preparation Kits following the manufacturer’s recommendations (Illumina, Ipswich, MA, USA). We had rRNA depletion performed during library preparation so as to preserve non-polyadenylated prokaryotic reads, enabling quantification of gene expression for both eukaryotes and prokaryotes. We distributed RNA samples from the same treatment across different lanes to preclude sequencing lanes as a batch effect. See Methods S2 for more details regarding RNA extraction, library preparation, and sequencing methods. Raw read data is available through the Sequence Read Archive under BioProjectID PRJNA1090526.

### Dual RNA-seq data analysis and gene expression quantification

After sequencing, we used the default settings of Trimmomatic (Bolger *et al*., 2014) to remove sequencing adapters from reads and discard poor quality reads. We used Sortmerna (Kopylova *et al*., 2012) and the rRNA sequences of all three species in the study system in the SILVA database (Glöckner *et al*., 2017; Pruesse *et al*., 2007; Quast *et al*., 2013; Yilmaz *et al*., 2014, p. 2) to remove rRNA reads that were not removed by rRNA depletion. Concatenated genomes were created by concatenating the reference genomes (Methods S3) for the one or two organisms present in each sample. The remaining non-rRNA and high quality reads were aligned with default settings to the appropriate concatenated genomes for each sample (Espindula *et al*., 2020) using HISAT2 (Kim *et al*., 2019). Reads with alignment quality scores <30 and whose paired read aligned to a separate chromosome or scaffold were discarded. The remaining alignments were used with the union method from HTSeq-count (Putri *et al*., 2022) to quantify gene expression of all genomic features not annotated as repeat regions, rRNA, or tRNA in concatenated reference genome annotations.

We found our gene expression quantification data to be suitable for downstream analyses (Chung *et al*., 2021; Love *et al*., 2014): host gene expression quantification data segregated on a principal component analysis by sample type (Figure S**1**) and the percentage of genes with a coefficient of variation within at least one treatment that was greater than across all samples never exceeded 20% (Table S**5**), indicating biological variation was greater than technical variation for most genes. See Methods S3 for greater details regarding reference genomes and software.

### Differential expression analyses

We considered genes with an aligned read count of >1 RPKM within its genome as expressed (Hebenstreit *et al*., 2011) and applied this categorization across all samples. To identify differentially expressed genes (DEGs), we used DESeq2 (Love *et al*., 2014) with a likelihood ratio test and Benjamini–Hochberg multiple test correction on all genes expressed in a given organ. We used several DESeq2 models to test for differential expression across different sample types. See Methods S4 for details about the models used with DESeq2.

### Gene ontology overrepresentation analyses

We used Gene Ontology overrepresentation analyses (GOORA) of gene lists generated from our differential expression analyses to formulate hypotheses about the biological processes that may be different across our treatments. For each *M. truncatula* organ in our analysis, we created a custom database of the genes that were expressed in that organ and their corresponding Gene Ontology (GO) annotations from the *M. truncatula* genome annotation. We used the default settings (p value 0.05, q value 0.2, Benjamini–Hochberg correction) of the enrichGO function from the clusterProfile package in R (Wu *et al*., 2021) against the aforementioned databases to identify over represented GO terms in the gene lists generated by our differential expression analyses.

### Phenotypic effects of rhizobia strains on host fitness and nematode colonization

We recapitulated the experiment as described above with an additional 76 plants (19 per treatment group). Upon harvesting, we recorded and removed a number of nodules from a subset of 7-8 plants from each treatment for a separate experiment. We separated root and shoot systems and dried the shoot systems for >48 hours before weighing them. Root systems were frozen for later processing. We scanned root systems at 1200 dpi following York (2023). After scanning all roots, root systems from plants infected with nematodes were stained with red dye following the procedure outlined by Thies *et al*. (2002) to make nematode galls more apparent and then scanned them. We manually inspected dyed and undyed images and manually counted nodules and nematode galls. RootPainter (Smith *et al*., 2022), a deep learning tool designed to use convolutional neural networks to segment root images, was used to segment the undyed scanned root images and RhizoVision Explorer (Seethepalli and York, 2020; Seethepalli *et al*., 2021) was then used to quantify root architecture from segmented images. Specifically, we examined above ground biomass as a proxy for host fitness (Younginger *et al*., 2017), root biomass, total number of branch points and root tips, network area, root system perimeter and average, maximum, and median root diameter as indications of host nutrient acquisition strategy (van der Bom *et al*., 2020).

We used linear mixed-effects models and analysis of variance in R to examine relationships between traits and rhizobia strain and nematode effects. We fit our models to the data using the lme4 (Bates *et al.,* 2015) and used the Satterthwaite’s degrees of freedom method executed by the lmerTest package (Kuznetsova *et al*., 2017) to test for significance. We used the DHARMα package (Hartig, 2022) to check all of our models for outliers, normality, and homoscedasticity after fitting them to our data.

## Results

### Mutualist strain impacts the expression of host genes related to defense, development, and pathogenesis at the site of host-parasite interactions

Plant host gene expression in nematode galls was different in hosts inoculated with different rhizobia strains. Out of the 51,653 examined genes in the *M. truncatula* genome, 23,069 genes (44.661%) were expressed. Of these genes, 285 genes (1.24%) were differentially expressed (DE) across hosts with different rhizobia strains. Of these DEGs, the majority (208 genes; 73.0%) had higher expression in hosts inoculated with WSM1022 than hosts inoculated with USDA1021 (Figure **2a**,**c**).

**Figure 2:**
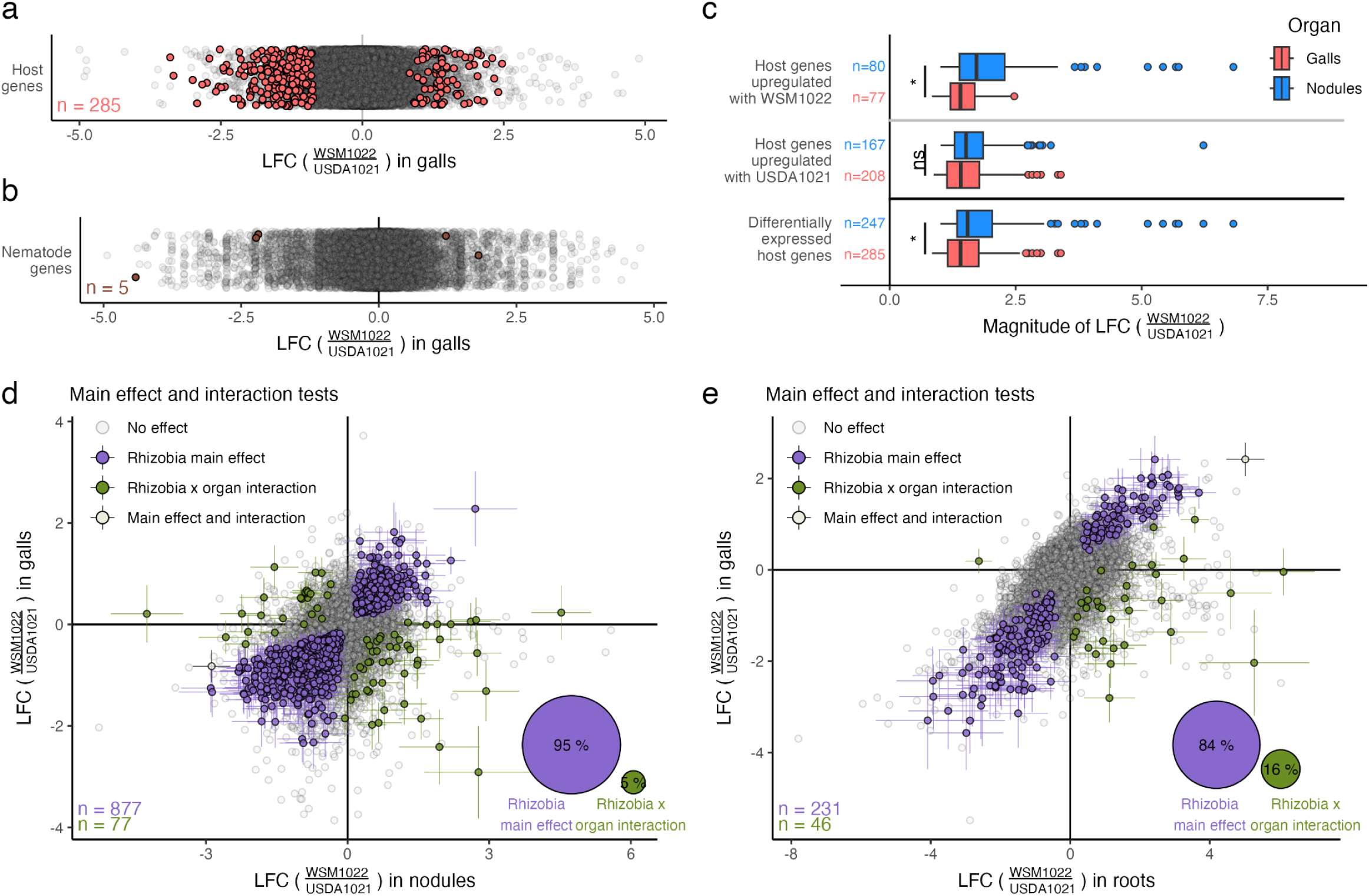
Rhizobia strain influences gene expression at the site of host-parasite interactions with largely similar responses in gene expression occurring in nematode galls and nodules and in nematode galls and roots. Log fold change (LFC) and standard error values were calculated using DESeq2. LFC values are the logarithm base 2 of the ratio of normalized mean expression in hosts inoculated with USDA1021 to that in hosts inoculated with WSM1022. Genes with significant log fold changes determined by likelihood ratio tests performed by DESeq2 are indicated with color and also with standard error bars in D and E. a) 261 host genes were differentially expressed (DE) in nematode galls across hosts with different rhizobia strains. b) Six nematode genes were DE across hosts with different rhizobia strains. c) Similar numbers of genes were DE in nematode galls and nodules across hosts with different rhizobia strains, but the magnitude of differential expression was higher in nodules. Differences in magnitude were tested using the optimal pooled t-test (Guo and Yuan, 2017) (‘n.s.’ indicates no significance, * indicates p-value < 0.05). d) Rhizobial differences produced largely similar responses in host gene expression in nodules and nematode galls. 712 genes showed a similar response to rhizobia strain in nodules and nematode galls (main effect) while 136 showed different responses in the two organs (interaction effect). e) Rhizobial differences produced largely similar responses in host gene expression in roots and nematode galls. 191 genes showed a similar response to rhizobia strain in roots and nematode galls (main effect) while 31 showed different responses in the two organs (interaction effect).

GOORA showed that annotations associated with diverse biological processes are overrepresented in all 285 host DEGs that responded to rhizobial strain differences in nematode galls (Dataset S6). Notably, genes annotated as related to carbohydrate metabolism were overrepresented. Functions related to the plant hormone abscisic acid, which coordinates plant responses to biotic and abiotic stress (Cao *et al*., 2011), were also overrepresented. Functions related to metabolism of cell wall components such as xylan, hemicellulose, arabinan, and pectin, and cell wall organization and biogenesis were also overrepresented. Changes in plant cell walls are both a symptom of and defense response against root knot nematode infection (Caillaud *et al*., 2008; Przybylska & Obrępalska-Stęplowska, 2020). Phosphoprotein regulation was also overrepresented in the host gene expression changes (Table S**2a**; see Dataset S6 for full details of GOORA results).

GOORA produced similar results when performed with only the DEGs in nematode galls that were upregulated in hosts inoculated with USDA1021. In contrast, within the DEGs in nematode galls that were upregulated in hosts inoculated with WSM1022, galacturan 1,4-alpha-galacturonidase activity (GO:0047911) and carbohydrate metabolic process (GO:0005975) were the only overrepresented annotations (Dataset S6). The activity of galacturan 1,4-alpha-galacturonidase degrades pectin in plant cell walls (Hasegawa and Nagel, 1968). Galacturan 1,4-alpha-galacturonidase enzymes are also secreted by root knot nematodes into cells at the infection site and pectin de-esterification and degradation is critical for successful root knot nematode infection (Jagdale *et al*., 2021; Wieczorek *et al*., 2014). Pectin metabolism and homeostasis affects many developmental pathways in plants (Guo *et al*., 2022; Wolf *et al*., 2009). Together these ORA results suggest the possibility that host carbohydrate metabolism, developmental processes, defense responses, or host pathogenesis are potentially altered in nematode galls when hosts are colonized by these two different rhizobia strains.

### Mutualist strain has little impact on global parasite gene expression but does alter a putative parasitism gene

Nematode gene expression showed little response to differences in the rhizobia strains resident in host nodules. Out of the 14,419 genes in the *M. hapla* genome, 12,880 genes (89.326%) were expressed under the aforementioned expression criteria. Of these genes, only five (0.04%) were DEGs (Figure **2b**, Table S**1a**).

Many parasitic nematode effector proteins localize to the nucleus where they modulate host gene expression, so we investigated the predicted protein products of the five differentially expressed genes for secretory signal sequences using SignalP 6.0 (Teufel *et al*., 2022) and for nuclear localization signals using NLStradamus (Nguyen Ba *et al*., 2009). Four of the five DEGs (MhA1_Contig1073.frz3.gene2, MhA1_Contig1386.frz3.gene15, MhA1_Contig253.frz3.gene16, and MhA1_Contig953.frz3.gene1) were predicted to code for secreted proteins by Signal 6.0. One of these four genes (MhA1_Contig1073.frz3.gene2) is also predicted to contain a nuclear localization signal sequence by NLStradamus. The protein product of the predicted cylicin-2 gene (MhA1_Contig253.frz3.gene16) also shows significant sequence similarity (coverage of 95% and percent identity 81.13%) to Minc00331 in the closely related *Meloidogyne incognita* which is presumed to be a parasitism gene as it is upregulated during parasitic life stages (Nguyen *et al*., 2018).

### Mutualist strain variation elicits similar responses in nodules and nematode galls

Out of the 26,875 *M. truncatula* genes expressed in all three organ types of nematode-infected hosts (52.030% of the genome), 9 genes showed a similar response to rhizobia strain across all organ types (Table S**1c**) and 705 genes (2.62% of expressed genes) showed a rhizobia strain by organ interaction effect. Notably, one gene that responded to rhizobia strain across all organ types (MtrunA17_Chr8g0364961/MtTPS9) and showed higher expression in hosts colonized by USDA1021 has been shown to be upregulated in response to abiotic stress and may mediate jasmonic acid response (Song *et al*., 2021). This indicates that the response to different rhizobia strains varies across organs and that very few host genes are responding similarly in all organs to intraspecific genetic variation in rhizobia.

Few genes are responding to differences in mutualist genotypes in similar ways across roots, nodules, and nematode galls, but similar responses can be observed across pairs of organs. Out of the 25,732 *M. truncatula* genes expressed in both symbiotic structures – nodules and nematode galls – 877 genes showed a similar response and 77 genes showed a different response to mutualist strain (Figure **2d**), indicating that the majority of genes that respond to different mutualist strains in nematode galls and nodules (81%) are responding in similar ways. Despite the lack of systemic response to different rhizobia strains across all organs in this study, symbiotic organs are responding similarly to differences in rhizobial strain. GOORA of the genes responding similarly in nematode galls and nodules suggests that processes associated with carbohydrate metabolism, cell walls, signal transduction, phosphorylation, and rhythmic/circadian processes may be responding in similar ways in the two symbiotic organs (Dataset S6). When changes in nematode galls were compared to changes in roots, we found that out of the 25,086 *M. truncatula* genes expressed in nematode galls and roots, 231 genes showed a similar response and 46 genes showed a different response to rhizobial strain across the two organ types (Figure **2e**), indicating that the majority of genes in nematode galls and roots that respond to rhizobial differences (83%) are responding to mutualist strain in similar ways in the two organs.

### Parasite infection impacts gene expression at the site of host-mutualist interactions in both strains of rhizobia

Parasite infection caused little change in gene expression in rhizobia. WSM1022 showed no DEGs between infected and uninfected hosts and USDA1021 had only 13 DEGs (Figure **3b**, Table S**1b**). In contrast, the host showed a large transcriptomic response in nodules to parasite infection. In nodules with rhizobia strain USDA1021, 22,993 *M. truncatula* genes (46.514% of the genome) were expressed. Of the expressed genes, 1,463 host genes (6.363%) were differentially expressed and similar numbers were up-and down-regulated in response to parasite infection (806 and 657 genes respectively) (Figure **3a**). Of these responding genes, 136 of them (9.30%) were conditionally expressed with 52 expressed only when parasites are present and 84 expressed only when parasites were absent. GOORA show that diverse processes are overrepresented in all 1,463 responsive host genes, but notably, terms related to host-mutualist interactions were overrepresented such as carbohydrate metabolic processes including oligosaccharide and glycogen metabolisms, energy reserve metabolism, defense response, transmembrane transport, structural constituents of cell walls, iron ion binding, and tripeptidyl-peptidase activity (Tables S**2b**,**c**,**d**; Dataset S**6**).

**Figure 3:**
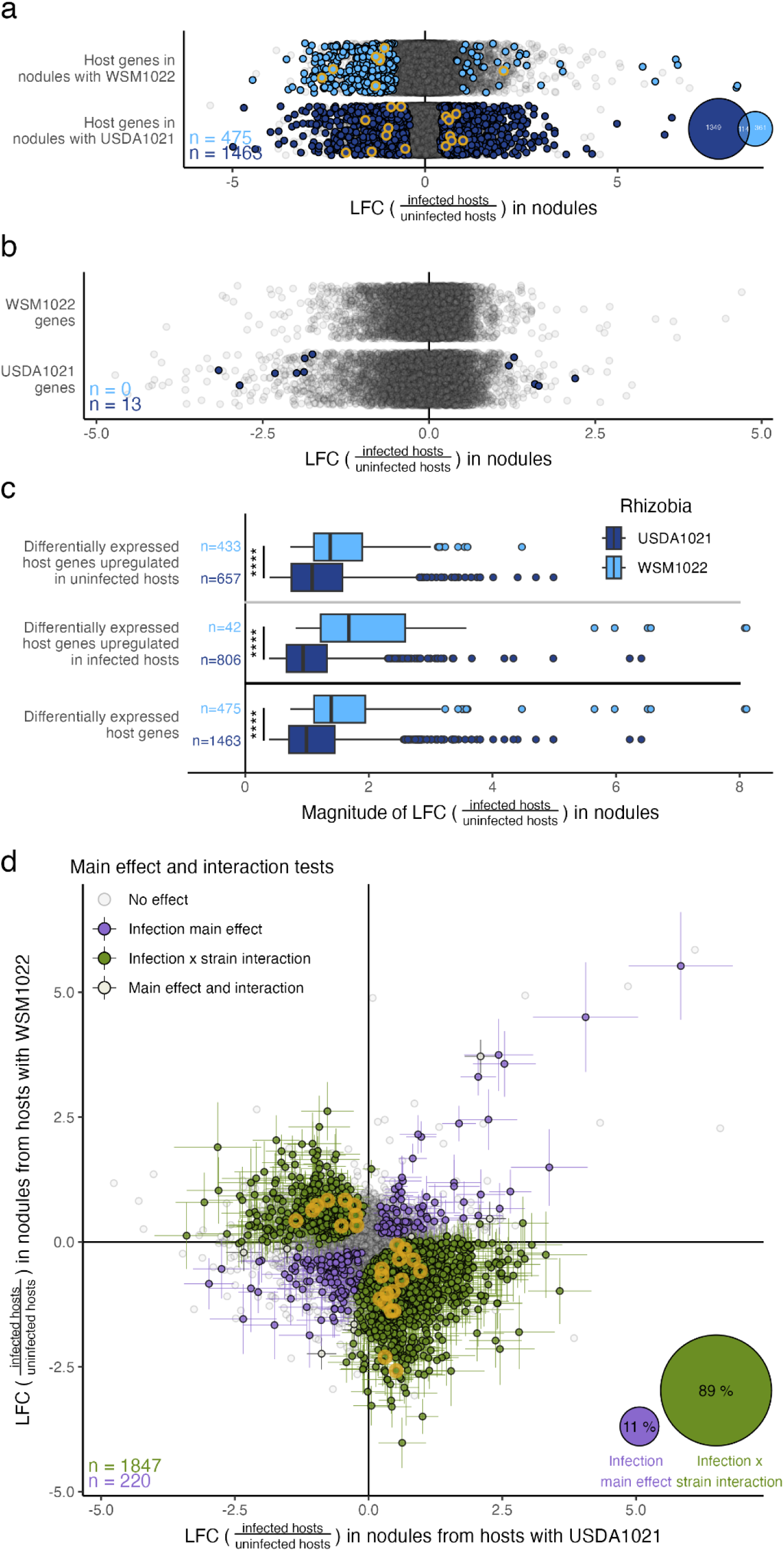
Nematode infection influences gene expression at the site of host-mutualist interactions in a largely strain specific manner. Log fold change (LFC) and standard error values were calculated using DESeq2. LFC values are the logarithm base 2 of the ratio of normalized mean expression in infected hosts to that in uninfected hosts. Genes with significant log fold changes determined by likelihood ratio tests performed by DESeq2 are indicated with color and also with standard error bars in d. All genes that were differentially expressed (DE) and functionally verified as critical to the functioning of the mutualism by Roy *et al*. (2019) have been circled in gold. a) 386 host genes were DE in response to nematode infection in nodules with WSM1022 and 1,428 were DE in nodules with USDA1021. b) 13 genes from USDA1021 and no genes from WSM1022 were DE in response to host nematode infection. c) The magnitude of host differential expression was higher in nodules with WSM1022. Differences in magnitude were tested using the optimal pooled t-test (B. Guo & Yuan, 2017) (‘n.s.’ indicates no significance, * indicates p-value < 0.05, ** indicates p-value < 0.01, *** indicates p-value < 0.001, **** indicates p-value < 0.0001). d) 1,632 host genes responded to parasite infection differently in nodules with different rhizobia strains (interaction effect) whereas only 193 genes responded similarly to parasite infection regardless of rhizobia strain (main effect).

In nodules with WSM1022, 22,729 *M. truncatula* genes (44.003% of the genome) were expressed. Of the expressed genes, 475 genes (2.09%) were differentially expressed and the majority (433 genes; 91.2%) were downregulated in response to parasite infection (Figure **3a**). Of the responding genes, 25 of them (5.8%) were conditionally expressed with 6 of them expressed only in the presence of parasites and 19 of them expressed only in the absence of parasites. GOORA suggests that parasite infection may impact the expression of host defenses as processes related to defense were overrepresented in all 475 responsive host genes as annotations related to defense response, regulation of salicylic acid biosynthesis, response to chitin, stress response, signal transduction, and cell wall related terms were overrepresented (Table S**2e**; Dataset S**6**).

### Parasite infection impacts host gene expression at the site of host-mutualist interactions in a largely mutualist strain specific manner

Out of the 23,565 host genes expressed across all nodule samples, 230 genes (0.976%) responded similarly to parasite infection in nodules regardless of mutualist strain. On the other hand, 1,847 of the expressed genes (7.838%) showed a strain specific response to parasite infection. This indicates that the majority of genes responding to parasite infection in nodules (88.93%) respond in different ways to mutualist genotype (Figure **3d**). Notably many of these genes responded in opposite ways when different rhizobia were resident in the nodule. Of these genes showing a strain-specific response, 645 genes (34.9%) showed conditional expression in which the gene was not expressed in at least one treatment.

GOORA showed that processes potentially involved in host-mutualist interactions are overrepresented in the host genes showing a strain specific response to parasite infection.

Notably, terms related to stress and defense responses, response to other organisms, cell communication and signal transduction, protein phosphorylation, autophosphorylation, and kinase activity, carbohydrate and energy reserve metabolism of various types as well as the transferase or hydrolyzing activity of glycosyls or sugars, and various processes related to components or organization of the cell wall were overrepresented. Additionally, GO terms related to cellular context showed that the genes showing a strain specific response were overrepresented for anchored components of the plasma membrane, plasmodesma, and cell walls (Table S**2f**; see Dataset S6). As many of these functions are important for plant-rhizobia interactions, especially carbohydrate metabolism (Udvardi and Poole, 2013) and defense responses (Cao *et al*., 2017; Gourion *et al*., 2015), this suggests that the strain-specific effect of parasites on host gene expression in nodules may influence host-mutualist interactions.

The examination of individual genes of interest can underscore the potential functional import of the strain-specific response in the transcriptomic data. The leghemoglobin gene MtrunA17_Chr5g0427351, a critical component of creating an environment within nodules that is conducive to nitrogen fixation (Larrainzar *et al*., 2020), responded to parasite infection in a strain-specific manner (Dataset S3). Roy *et al*. (2019) compiled a list of 207 genes who have been functionally verified as integral to the proper functioning of the *M. truncatula*-*E. meliloti* mutualism. Out of these 207 genes, 25 genes were differentially expressed in nodules in a strain specific manner (interaction effect) while none of these genes showed a consistent response to nematode infection (main effect) across hosts regardless of rhizobia strain (Figure 3d).

### Rhizobia strain had no effect on root architecture while nematode infected plants had thicker and shorter roots with fewer root tips

A number of the root architecture traits were highly correlated with one another (Figure S4; S5). Linear mixed models showed that nematode infection increased median root diameter (χ^2^_df=1_ = 4.26, p = 0.039) and average root diameter (χ^2^_df=1_ = 7.14, p = 0.008) and decreased root perimeter (χ^2^_df=1_ = 5.61, p = 0.018), total root length (χ^2^_df=1_ = 4.66, p = 0.031) and number of root tips (χ^2^_df=1_ = 6.28, p = 0.012) (Figure S6). However, host plants colonized by different rhizobia strains showed no difference in root architecture traits. The number of branch points, maximum root diameter, and network area were all unaffected by rhizobia strain and nematode infection and our models showed no interaction effect of rhizobia strain and nematode infection for any root architecture traits (Figure S6).

### Neither nematode infection nor rhizobia strain altered aboveground biomass, a proxy for host fitness

Plants that formed more nodules had significantly higher above ground biomass (r = 0.48, p <0.001; Figure 4), although there was appreciable scatter in this relationship for plants with intermediate nodule numbers, confirming that both strains of rhizobia confer benefits to their host plant., However, rhizobia strain was not a good predictor of above ground biomass (Figure 4d), indicating that the two strains did not differ in the benefits they provide their host. Neither nematode infection status nor parasite load (nematode gall count) affected aboveground biomass (Figure 4e), even when correcting for nodule count or root biomass. Together this suggests that, in concordance with other published data (Wood *et al.,* 2018), nematode infection status and nematode infection severity did not affect above ground biomass but nodulation rate did.

**Figure 4:**
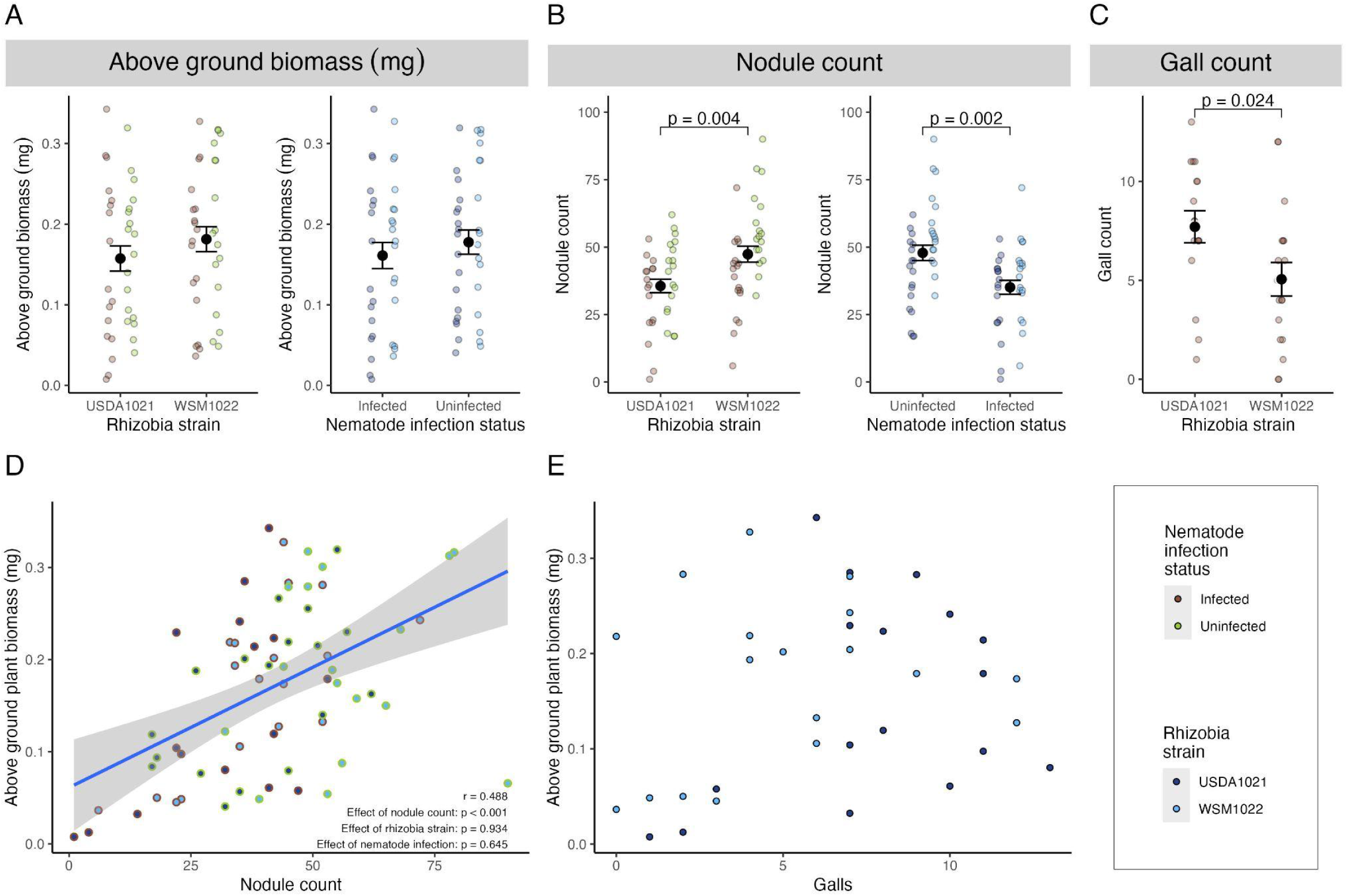
Rhizobia strain and nematode infection status could not explain variation in above ground biomass, but nodule count could. a) The above ground biomass of host plants did not differ across rhizobia strain or nematode infection status. b) Plants differed in their nodules counts across both rhizobial strain and nematode infection status. Infected plants had fewer nodules than uninfected plants, and plants inoculated with USDA1021 had fewer nodules than plants inoculated with WSM1022. c) Nematode-infected plants had different number of nematode galls depending on the rhizobia strain they were inoculated with. Plants inoculated with WSM1022 had fewer nematode galls than plants inoculated with USDA1021. d) Nodule count was a good predictor of above ground biomass but neither rhizobia strain nor nematode infection status changed the relationship between nodule count and host plant above ground biomass. e) Nematode gall count was not a good predictor of above ground biomass.

### Nematode infection status impacted nodule count and rhizobia strain impacted nematode gall count

As expected, plants with larger root systems had more rhizobia nodules and nematode galls (Figure S4; S5). Nodule and nematode gall counts were positively correlated (r = 0.42), and a linear mixed model verified rhizobia colonization rates is a good predictor of nematode colonization rates (χ^2^_df=1_ = 10.44, p = 0.001; Figure S6). This held true even when correcting for root volume (χ^2^_df=1_ = 3.86, p = 0.049), indicating that the relationship between mutualist and parasite colonization is not merely a byproduct of overall plant size. Plants inoculated with WSM1022 had a lower mean nematode gall count than plants inoculated with USDA1022 (χ^2^_df=1_ = 5.11, p = 0.024; Figure 4c), even after correcting for root biomass (χ^2^_df=1_ = 5.88, p = 0.015). Nematode decreased the number of nodules (χ^2^_df=1_ = 9.56, p = 0.002; Figure 4b) even when corrected for by root biomass (χ^2^_df=1_ = 12.96, p < 0.001) and WSM1022 produced higher nodule counts than USDA1021 (χ^2^_df=1_ = 8.14, p = 0.004; Figure 4b) even when corrected for by root biomass (χ^2^_df=1_ = 11.14, p < 0.001). In summary, we saw that rhizobia strain had an effect on gall counts and nematode infection decreased nodule counts.

## Discussion

Here we showed that intraspecific genetic variation in a major microbial mutualist impacts host-parasite interactions at the phenotypic and transcriptomic levels. Rhizobia strains affected their host’s parasite load (Figure 4c). At the mature site of parasite infection (nematode galls), hundreds of genes were differentially expressed across plant hosts that were inoculated with one of two different strains of mutualistic rhizobia. At the mature site of mutualist colonization (rhizobial nodules), the effect of parasite infection on host gene expression depended on the resident mutualist strain. Our results suggest that microbial indirect genetic effects play an underappreciated role in structuring their host’s interactions with other co-colonizing microorganisms at multiple stages of host-symbiont interaction.

### Microbial mutualists influence their host’s parasite load and gene expression at the site of parasite infection

Host plants inoculated with different strains of mutualistic rhizobia showed different numbers of nematode galls (Figure 4c) indicating that microbial mutualist genetic variation can affect the severity of parasite infection. This finding is in line with a variety of other studies that highlight the importance of microbial mutualists on traits that mediate other species interactions with hosts (Lange *et al*., 2023; Osborne *et al*., 2009; Parker *et al*., 2017). The difference in parasite infection severity that we observed was based on a count of mature parasitic structures and therefore could arise from differences in the speed at which parasite infection develops or differences in colonization rates due to varying host defenses or susceptibility. While we observed differences in parasite infection severity, host plant fitness proxies were not impacted by nematode infection (Figure 4). While this might suggest that nematodes were not acting parasitically in the present experiment, in past work they have been shown to reduce different fitness proxies than what was quantified here (Wood *et al*., 2018). Furthermore, under more realistic conditions where hosts may be exposed to more variable environmental conditions and resource availability, fitness consequences may be more apparent.

Strain-level variation in a microbial mutualist was a determinant of gene expression in parasite-infected hosts. This effect manifested in two ways. First, rhizobia had a comparable effect on host gene expression in nematode galls as in rhizobia nodules. The number of rhizobia-responsive plant genes in nematode galls was greater than in nodules, despite the magnitude of differential expression being slightly larger in nodules (Figure **2c**). Second, rhizobia altered the effect that parasite infection had on host gene expression. Hundreds of genes that were upregulated in response to parasite infection when the host was inoculated with one rhizobial strain were downregulated in response to parasite infection when inoculated with another strain (Figure **3c**).

Our results show that microbial effects on host gene expression extend to other species interactions their host engaged in. Furthermore, microbial effects on a host’s third-party interactions can rival their direct effect on hosts at their site of colonization. These results are consistent with and build on the growing body of evidence that microorganisms can strongly influence gene expression in their hosts: microbial modulation of host gene expression has been demonstrated in terrestrial and marine mutualisms (Burghardt *et al*., 2017; Heath *et al*., 2012; Palakurty *et al*., 2018; Rodriguez-Lanetty *et al*., 2006), in animal and plant pathogens (Apidianakis *et al*., 2005; Stringlis *et al*., 2018), and in the primate microbiome (Grieneisen *et al*., 2020).

### The consequences of cascading effects of mutualist strain variation

Simple models of selection on mutualisms predict that selection should remove intraspecific variation in mutualism-related traits when, in reality, extensive intraspecific variation is a hallmark of many mutualism-related traits (Heath and Stinchcombe, 2014). This is true in the legume-rhizobia mutualism as rhizobia strains vary dramatically in the fitness benefit they provide their host (Heath, 2010). Similar patterns have been documented in mutualisms across the tree of life, from mycorrhizal fungi (Hart *et al*., 2013), to obligate pollinators (Jandér *et al*., 2012), to the bacterial symbionts of marine mussels (Ansorge *et al*., 2019). This disparity has motivated searches for mechanisms that maintain variation in mutualisms (Heath and Stinchcombe, 2014). Our experiment suggests that the mechanisms maintaining variation in mutualisms may have unanticipated and underappreciated consequences for the ecological communities in which mutualisms are embedded. However, predicting when and how intraspecific variation in mutualisms will cascade to impact other species interactions in host-associated communities requires understanding the mechanisms mediating crosstalk between microbial mutualists and other co-colonizing microorganisms.

### Potential mechanisms underlying mutualist strain effects on host-parasite interactions

Why might strain-level variation in a microbial mutualist have an effect on the host’s response to a parasite? We elaborate on three non-exclusive explanations below: a defense-centric, resource-centric, and development-centric model.

First, mutualist strains may elicit different defense responses in their host that may cascade to impact the expression of defense responses at the site of parasite infection. The establishment and maintenance of many mutualisms requires tightly coordinated control of the host’s immune response (Zipfel and Oldroyd, 2017). In the legume-rhizobia mutualism, suppression of the plant’s immune response via nod factors and other effectors orchestrate the initiation and progression of the mutualism (Cao *et al*., 2017; Gourion *et al*., 2015). Mature nodules have a suppressed immune response, and functional genetic experiments have established that maintaining a functional symbiosis necessitates a controlled and tightly coordinated immune response (Zhang *et al*., 2023).

Consistent with this defense-centric model, our GOORA analysis of nematode gall samples uncovered enrichment of defense-related functions in DEGs. Notably, functions related to the plant hormone abscisic acid were overrepresented. Abscisic acid coordinates biotic and abiotic stress responses (Cao *et al*., 2011), although its role in plant responses to nematode infection remains poorly understood (Gheysen and Mitchum, 2019). changes in plant cell walls are a defense response against root knot nematode infection (Caillaud *et al*., 2008; Przybylska & Obrępalska-Stęplowska, 2020) and functions related to the metabolism of cell wall constituents were also overrepresented. Thus our findings about parasite severity may also be explained by rhizobia strains prompting varying defense responses in host plants that result in different degrees of host susceptibility. It is also notable that WSM1022 is a unique strain of rhizobia in that it contains the Type III Secretion System, whereas many other strains of rhizobia do not. The Type III Secretion System plays a versatile role in modulating the legume immune response (Teulet *et al*., 2022), and could produce differential systemic immune responses that lead to the phenotypic differences we observed.

Second, mutualist strains may differ in the resources they provide or costs they impose on their host. If strains differ in their impact on the host’s resource budget, these differences could cascade to impact resource flux at the site of parasite infection or produce differences in the way mutualism resource fluxes respond to parasite infection. Intraspecific variation in the costs and benefits a mutualist provides is well documented in the legume-rhizobia mutualism (Heath, 2010; Calvert *et al*., 2023) and rhizobia strains vary in the nitrogen and benefits they provide their plant host (Batstone *et al*., 2017; Terpolilli *et al*., 2008). If the strains in our experiment negotiated different carbon-for-nitrogen exchange rates with their host, the resulting impact on the host’s resource budget could have driven differences in resource flux in nodules, nematode galls, or both, driving the transcriptomic differences we observed.

In support of this resource-centric model, our GOORA analyses suggest that rhizobia strain affected carbohydrate metabolism in nematode galls and nematode infection affected carbohydrate metabolism in nodules (Dataset S6). Carbohydrate metabolism plays a central function in the legume-rhizobia mutualism (Udvardi and Poole, 2013), and root knot nematodes siphon carbohydrates from the phloem, facilitated by active sugar transport into nematode galls (Zhou *et al*., 2023). Furthermore, the expression of leghemoglobin, a key regulator of the nitrogen-fixation reaction in rhizobia nodules (Larrainzar *et al*., 2020), exhibited a strain-specific response to parasite infection. This observation raises the intriguing hypothesis that rhizobia strains control the sensitivity of nitrogen fixation—the central function of the legume-rhizobia mutualism—to parasite infection. Resource budget differences may also account for our phenotypic data, as parasite access to host resources can influence parasite infection severity or progress (Schultz *et al*., 2013; Compson *et al*., 2011). Differences in resource exchange across different rhizobia strains may alter host resource budgets in ways that impact the nematode’s ability to siphon nutrients from its host, further impacting nematode infection severity, host defenses, or parasite pathogenesis.

Third, mutualist strains may differ in their impact on developmental processes. Symbiotic microbes are known to have profound effects on host development in both plants and animals on a systemic and local level (Fronk and Sachs, 2022; O’Brien *et al*., 2021a; Sommer and Bäckhed, 2013). Furthermore, resistance to infectious disease changes across ontogeny: juveniles are typically more susceptible to infection than adults (Ashby and Bruns, 2018; Izhar *et al*., 2020). If mutualist strains have different impacts on host systemic ontogeny and host ontogeny influences parasite resistance or infection trajectory, then hosts inoculated with different mutualists could be in different stages of infection when sampled at the same chronological age. On a local scale, mutualist strain differences may alter nematode gall or nodule development. Regardless of scale, in bulk RNAseq, differences in ontogeny can produce signals of differential expression if the ratio of different cell types in sequenced bulk organs differs across treatments of interest (Hunnicutt *et al*., 2022; Montgomery and Mank, 2016). Therefore, the differential expression that we observed may result from rhizobia strains influencing nematode gall, nodule, or systemic development such that cell composition in nematode galls or nodules differed across treatments. Developmental differences in hosts may also impact nematode infection severity as host ontogeny can influence parasite susceptibility and infection severity (Burr *et al*., 2022).

### Outstanding questions and future directions

Our results suggest several promising avenues of future research. First, what are the relative contributions of defense-, resource-, and development-centric models to crosstalk in host-associated communities? Our GOORA implicates both the defense-and resource-centric models in mediating transcriptional crosstalk between the rhizobia mutualism and nematode parasitism but is unable to implicate development-centric models. Similarly, our phenotypic data can be explained by all three models. Future experiments should independently manipulate host defense and resource flux to disentangle their independent contributions to crosstalk between co-infecting mutualists and parasites. Similarly, future experiments should be designed to examine different developmental stages or the contribution of different tissue types within symbiotic organs of the host-parasite interaction (Martinson *et al*., 2022) to examine how rhizobial strain effects occur in different developmental stages. Experiments that implement sequencing technologies that de-confound the gene expression profiles of different cells or tissues, such as spatial transcriptomics (Rao *et al*., 2021) or scRNAseq (Saliba *et al*., 2014), could also shed light on the contribution of development to differential gene expression and phenotypic differences. Beyond illuminating the organismal processes that underlie crosstalk in host-associated communities, discriminating between defense-, resource-, and development-centric models is important to understand how these communities may evolve, as different underlying mechanisms implicate different regimes of pleiotropy and crosstalk that may constrain or alter evolutionary potential and shape organismal traits and species interactions. For example, if crosstalk is primarily mediated by the host’s defense response, co-colonization may impact the evolution of genes and processes involved in host immunity. Alternatively, if crosstalk is primarily resource mediated, co-colonizers may influence the evolution of resource allocation and acquisition traits in their hosts.

Second, how dynamic are the indirect effects of rhizobia strains on host-nematode interactions? We observed transcriptomic differences in mature galls from host plants that were inoculated with different rhizobia strains. By quantifying host susceptibility at a single time point, we may have missed a dynamic process in which rhizobial strains have an indirect influence on host-parasite interactions that vary over time and development. Plant genotype, age of host plant, and the time since symbiont inoculation appear to interact to influence host susceptibility to nematodes (Yi, Unpublished data). Furthermore, previous studies have shown that host age at colonization and the synchronicity of nematode and rhizobia inoculation affect host susceptibility to nematodes (Burr *et al*., 2022). Future experiments that incorporate multiple timepoints or different host ages may be able to identify whether there are specific windows of time where rhizobial strains have stronger or weaker indirect effects on host-parasite interactions.

Third, why was the transcriptomic response to co-colonization attenuated in symbionts relative to hosts? And why were changes in host gene expression in symbiotic structures not mirrored by the resident symbiont? In contrast to host gene expression, mutualist gene expression was largely unaffected by parasite infection, and parasite gene expression was largely unaffected by mutualist strain differences. Only 0-0.2% of rhizobia and nematode genes responded to their co-colonizer, while 1-6% of plant genes responded to variation in one or both symbionts. This observation suggests that the effect of mutualist strain on host-parasite interactions is primarily driven by the host. One straightforward explanation is that direct contact between species drives a larger transcriptomic response than indirect interactions. Another possibility is that generalist symbionts — like root-knot nematodes (Castagnone-Sereno *et al*., 2013; Catella *et al*., 2022), and, to a lesser extent, *Ensifer* rhizobia (Harrison *et al*., 2018) — have evolved transcriptional robustness across a wide variety of host environments. As mentioned above, it is also possible that transcriptomic responses of symbionts may vary across developmental time or as a function of host age.

Fourth, when does crosstalk in host-associated communities arise from systemic versus symbiosis-specific host responses? A significant insight from our experiment is that mutualist effects on gene expression at the site of parasite infection were specific to symbiotic organs and not a byproduct of systemic changes in host gene expression. Rhizobia elicited a different transcriptomic response in nodules and nematode galls than in uncolonized root tissue (Figure **2**). One possible explanation is that symbiosis-specific responses dominate whenever there is substantial overlap in the function of mature symbiotic structures induced by different colonizing species. Despite rhizobia being a mutualistic prokaryote and nematodes being a parasitic eukaryote, there is substantial overlap in their functions (Fronk and Sachs, 2022): both involve nutrient exchange, manipulation of the host defense response, and modification of host cell development and differentiation (Koltai *et al*., 2001). Regardless of the mechanism, our results underscore the importance of tissue specificity when studying the transcriptomic basis of host-symbiont interactions (Koltai *et al*., 2001; Przybylska and Obrępalska-Stęplowska, 2020).

Finally, our study raises a final fundamental question: How much genetic variation for traits mediating species interactions (e.g., parasite resistance and mutualism function) resides in species not directly involved in the interaction itself? Our results indicate that indirect genetic effects are important and overlooked coordinators of species interactions within multispecies contexts. The concept of indirect genetic effects—when one organism’s genes influence another’s phenotype—was originally formalized in the context of intraspecific interactions like aggression or maternal effects (Moore *et al*., 1997; Wolf *et al*., 1998). Recently, this framework was formally extended to interactions between species (De Lisle *et al*., 2022). However, this field primarily remains focused on pairwise species interactions: for example, how host genes influence parasite virulence or the benefit derived from a mutualistic symbiont. A significant insight from our study is that indirect genetic effects in species interactions extend to third parties. Given the potential that microbial communities have for impacting selection and overall health of their hosts (Lau and Lennon, 2011), future work should explore the role of intraspecific variation in microbial symbionts in shaping the structure and function of host-associated communities and its consequences for host health and fitness.

## Supporting Information

The following Supporting Information is available for this article:

**Dataset S1**: A comma-separated values file containing gene expression quantification data output from HTSeq-union for all genes in all organisms.

**Dataset S2**: A comma-separated values file containing gene expression status for all genes in all samples.

**Dataset S3**: A comma-separated values file containing the results of all DESeq2 differential expression analyses for all *Medicago truncatula* genes.

**Dataset S4**: A comma-separated values file containing the results of all DESeq2 differential expression analyses for all *Ensifer meliloti* genes.

**Dataset S5**: A comma-separated values file containing the results of all DESeq2 differential expression analyses for all *Meloidogyne hapla* genes.

**Dataset S6**: A comma-separated values file containing all Gene Ontology overrepresentation analysis results.

**Figure S1:** Primary component analysis of *M. truncatula* gene expression quantification data showing segregation of datapoints by organ type.

**Figure S2:** Upset plot showing the intersection of *M. truncatula* gene lists produced by differential expression analyses comparing similar and dissimilar responses to rhizobial strain differences across different groups of organs.

**Figure S3:** Diagrams showing the expression and differential expression status of *M. truncatula* genes in different organ types.

**Figure S4:** Plot showing the correlation between root architecture traits and nodules counts across all samples.

**Figure S5:** Plot showing the correlation between root architecture traits and nodule and gall counts across all samples that were infected with nematodes.

**Figure S6:** Plot showing root architecture traits separated by nematode infection status and rhizobia strain.

**Table S1:** Gene lists of interest.

**Table S2**: Select Gene Ontology terms that were enriched in various gene lists produced by differential expression analyses.

**Table S3**: Constituents of the various fertilizer solutions used in this experiment.

**Table S4**: Symbiont treatment (nematode status and rhizobial strain) and host organ of origin for each sample.

**Table S5**: Parameters for the DESeq2 likelihood ratio tests performed for each model.

**Table S6**: Table showing the percentage of genes in each organ and treatment that had a coefficient of variation within the treatment that was greater than the coefficient of variation across all samples.

**Table S7**: Table showing total reads and percentage of reads that are filtered or aligned at various bioinformatic steps.

**Methods S1**: Sample preparation

**Methods S2**: RNA extraction and sequencing

**Methods S3**: RNA Quality Control and Library Preparation Performed by Genewiz Azenta Life Sciences

**Methods S4**: Software and reference genomes

**Methods S5**: Differential expression analysis

**Results S1:** Nodule cysteine-rich protein gene expression is impacted by parasite infection

