## Supporting Information for "Variation in a microbial mutualist has transcriptional and phenotypic consequences for host-parasite interactions"

[authors removed for anonymity]

**Table of Contents:**

| | |
|---|---|
| Dataset lists | pg 2 |
| Figure S1 | pg 4 |
| Figure S2 | pg 5 |
| Figure S3 | pg 6 |
| Figure S4 | pg 7 |
| Figure S5 | pg 8 |
| Figure S6 | pg 9 |
| Table S1 | pg 10 |
| Table S2 | pg 12 |
| Table S3 | pg 16 |
| Table S4 | pg 17 |
| Table S5 | pg 19 |
| Table S6 | pg 21 |
| Table S7 | pg 23 |
| Supplemental Methods (Methods S1-S4) | pg 24 |
| Supplemental Results (Results S1) | pg 28 |

All datasets and scripts to generate the datasets can be accessed at:
www.github.com/adbmar/MtruncDualRNASeq2024

All datasets are also available at:
doi:10.5061/dryad.b8gtht7mp

***Dataset S1:*** *Gene expression quantification*

A comma-separated values file containing gene expression quantification data output from HTSeq-union for all genes in all organisms.

***Dataset S2:*** *Gene expression status*

A comma-separated values file containing gene expression status for all genes in all organisms. Expression status is determined by a gene having >1 RPKM (reads per kilobase of gene length per million reads of an organism) in more than one sample of a given treatment. Column names are composed of the string "exp_" followed by an abbreviation of the organ type and the rhizobia strain treatment. The abbreviation "nodp" indicates nodules from hosts with inoculated with nematodes and the abbreviation "nodm" indicates nodules from hosts without nematode inoculation. "21" indicates rhizobial strain USDA1021 and "22" indicates rhizobial strain WSM1022. The abbreviation "gano" indicates the combination of galls and nodules, the abbreviation "garo" indicates the combination of galls and roots, and the abbreviation "noro" indicates the combination of nodules from hosts infected and roots.

***Dataset S3:*** *Differential expression analysis for Medicago truncatula*

A comma-separated values file containing the results of all DESeq2 differential expression analyses for *M. truncatula* genes. Column names are composed of a prefix and suffix separated by a period. Each prefix corresponds to one of the models detailed in Table **S3**. Suffixes are as follows: "baseMean" indicates the mean of normalized expression across all sample types; "ns_LFC" indicates the log fold change base 2 of the normalized expression across the contrast specified by the model in Table **S3**; "ns_SE" indicates the standard error of the log fold change estimate; "padj" indicates the p-value of the likelihood ratio test used to determine if the log fold change is significantly different from zero adjusted with false discovery rate correction. The final column entitled NCR indicates whether a gene is a nodule-specific cysteine-rich protein.

***Dataset 4:*** *Differential expression analysis for Ensifer meliloti*

A comma-separated values file containing the results of all DESeq2 differential expression analyses for all *E. meliloti* genes. Column naming convention follows the same pattern as described in Dataset 3.

***Dataset 5:*** *Differential expression analysis for Meloidogyne hapla*

A comma-separated values file containing the results of all DESeq2 differential expression analyses for all *M. hapla* genes. Column naming convention follows the same pattern as described in Dataset 3.

***Dataset 6:*** *Gene Ontology overrepresentation analysis*

A comma-separated values file containing all Gene Ontology overrepresentation analysis results. These results are the output of the enrichGO function in the clusterprofiler R package.

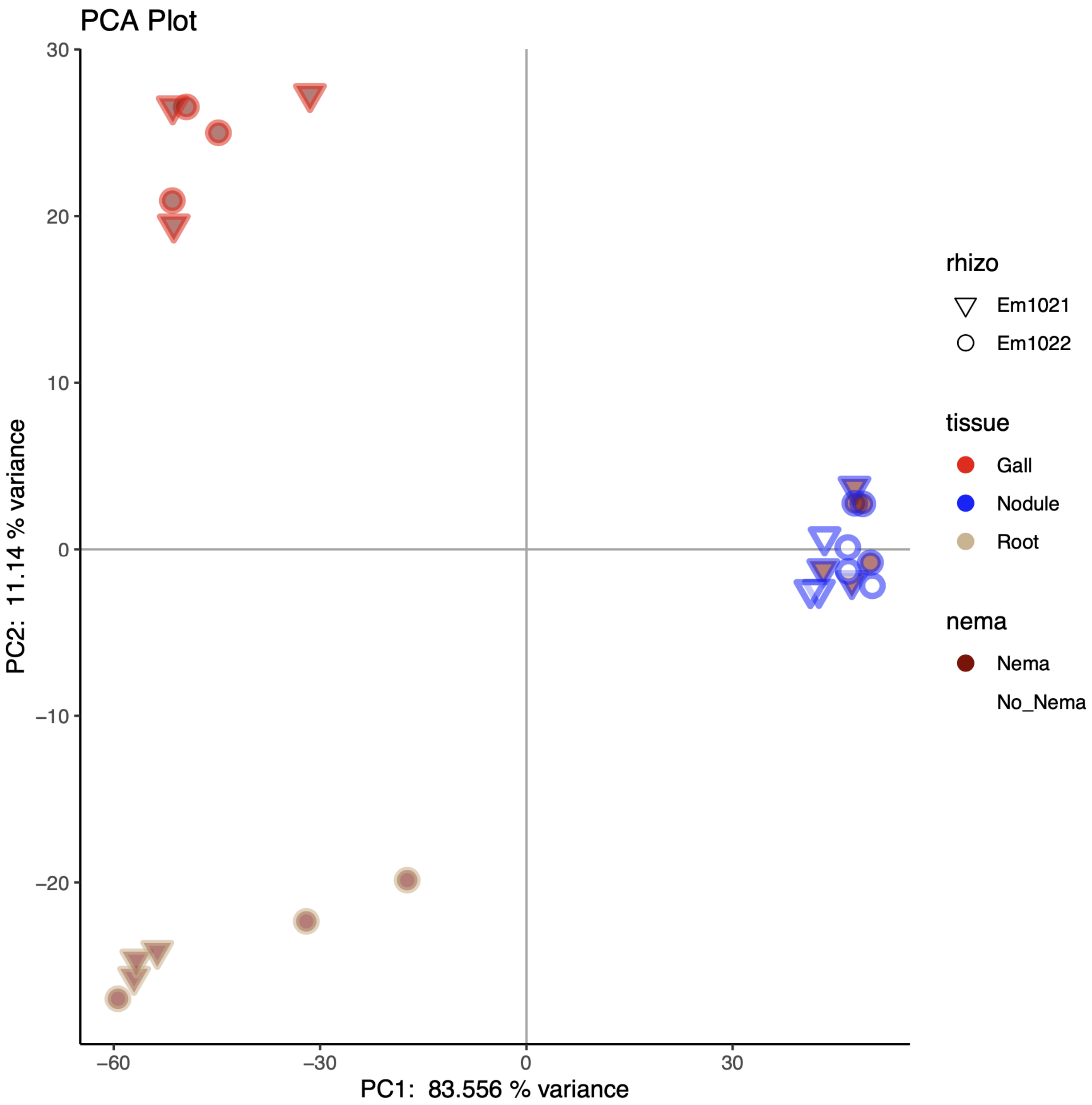

**Fig S1**: Primary component analysis (PCA) of gene expression quantification data of *Medicago truncatula* genes. Each point represents the PCA projection of the gene quantification data output from HTSeq-union for a single sample. Samples segregated by organ type as indicated by the clumps of similarly colored data points.

Similar or different responses to rhizobia strain across different sets of organs

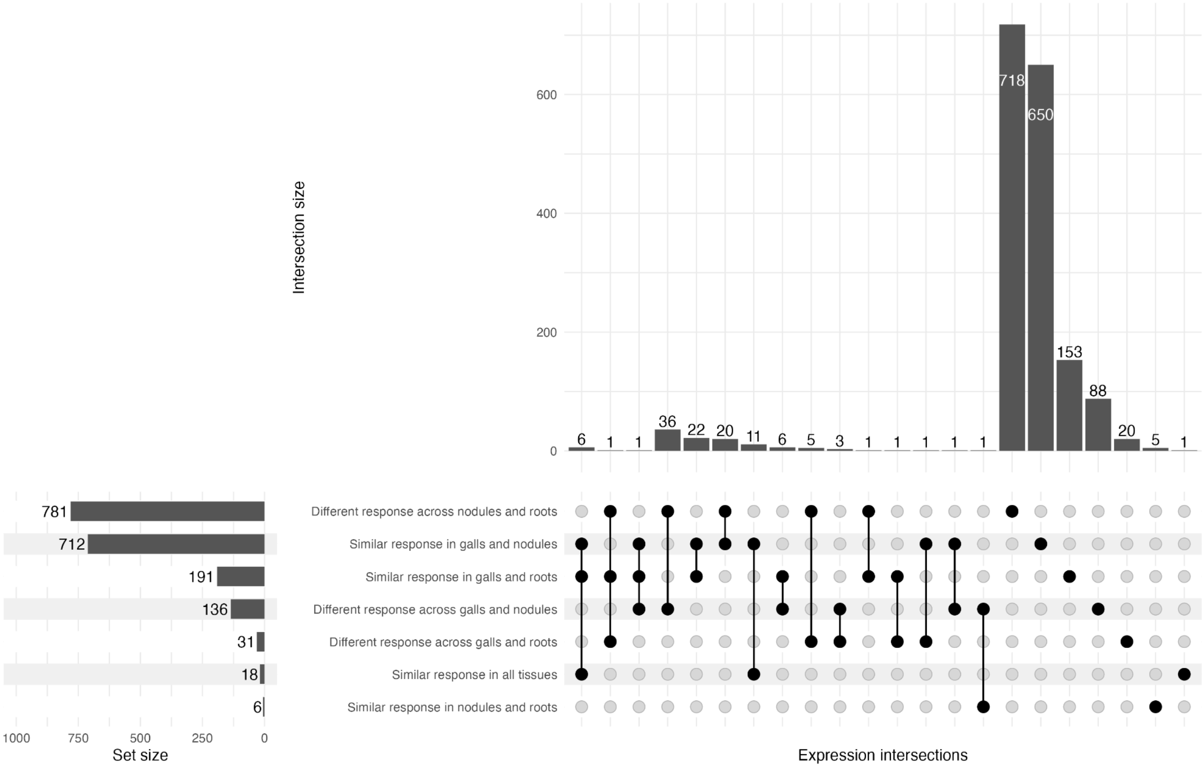

**Fig. S2**: Upset plot showing the intersection of *M. truncatula* gene lists produced by differential expression analyses comparing similar or different responses to rhizobial strain (statistically, a main effect of rhizobia strain or rhizobia x organ interaction respectively) across different groups of organs.

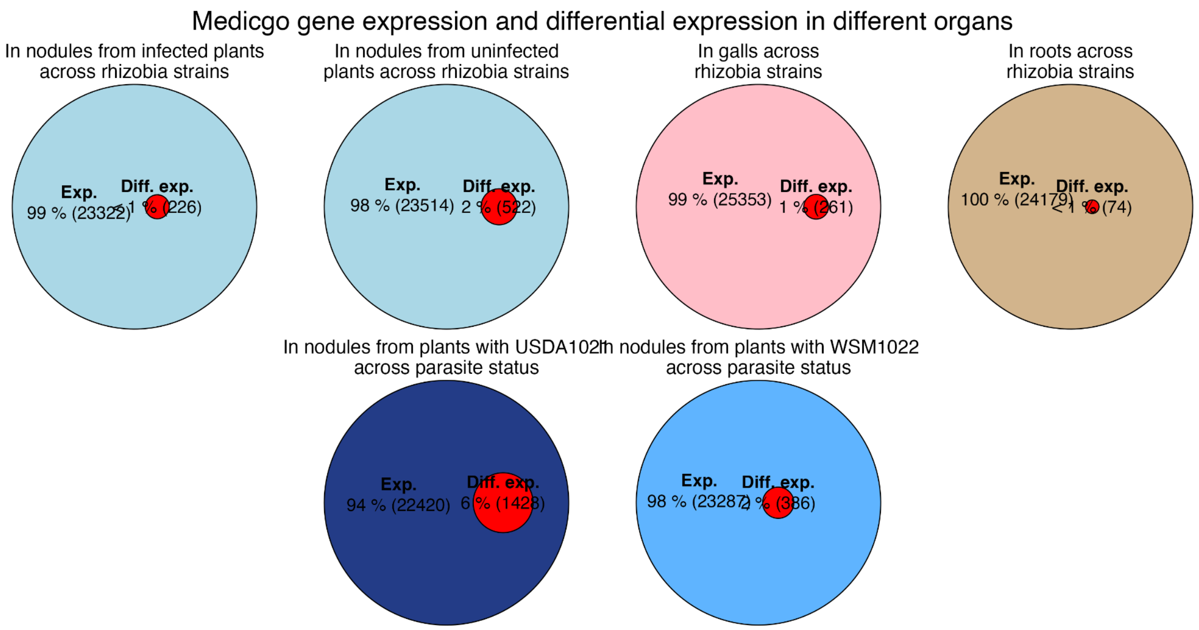

**Fig. S3**: Venn diagrams showing the number and percentage of expressed genes that were also differentially expressed in different organs and across different comparisons.

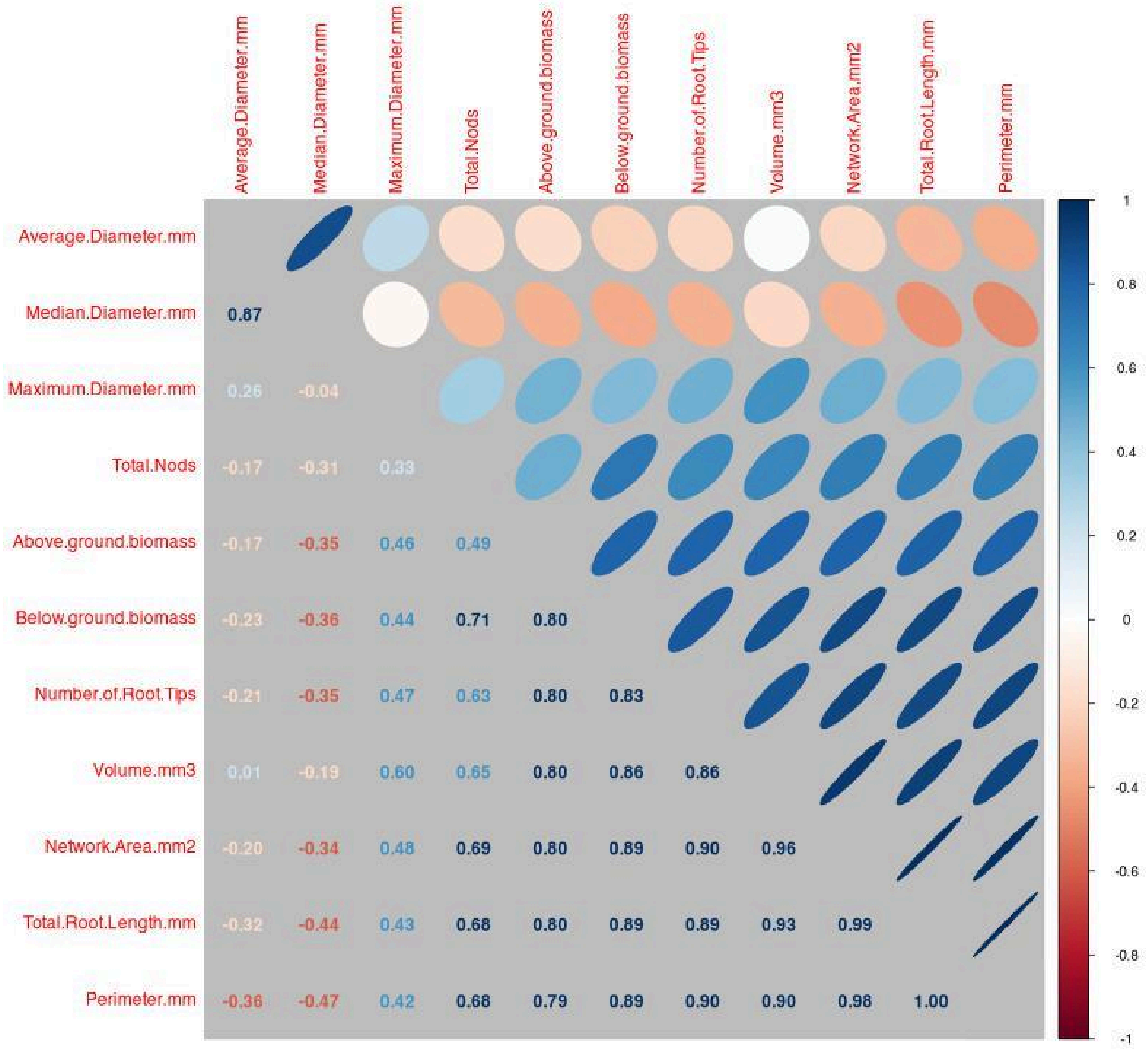

**Fig. S4**: Plot showing the correlation between root architecture traits and nodule counts across all samples.

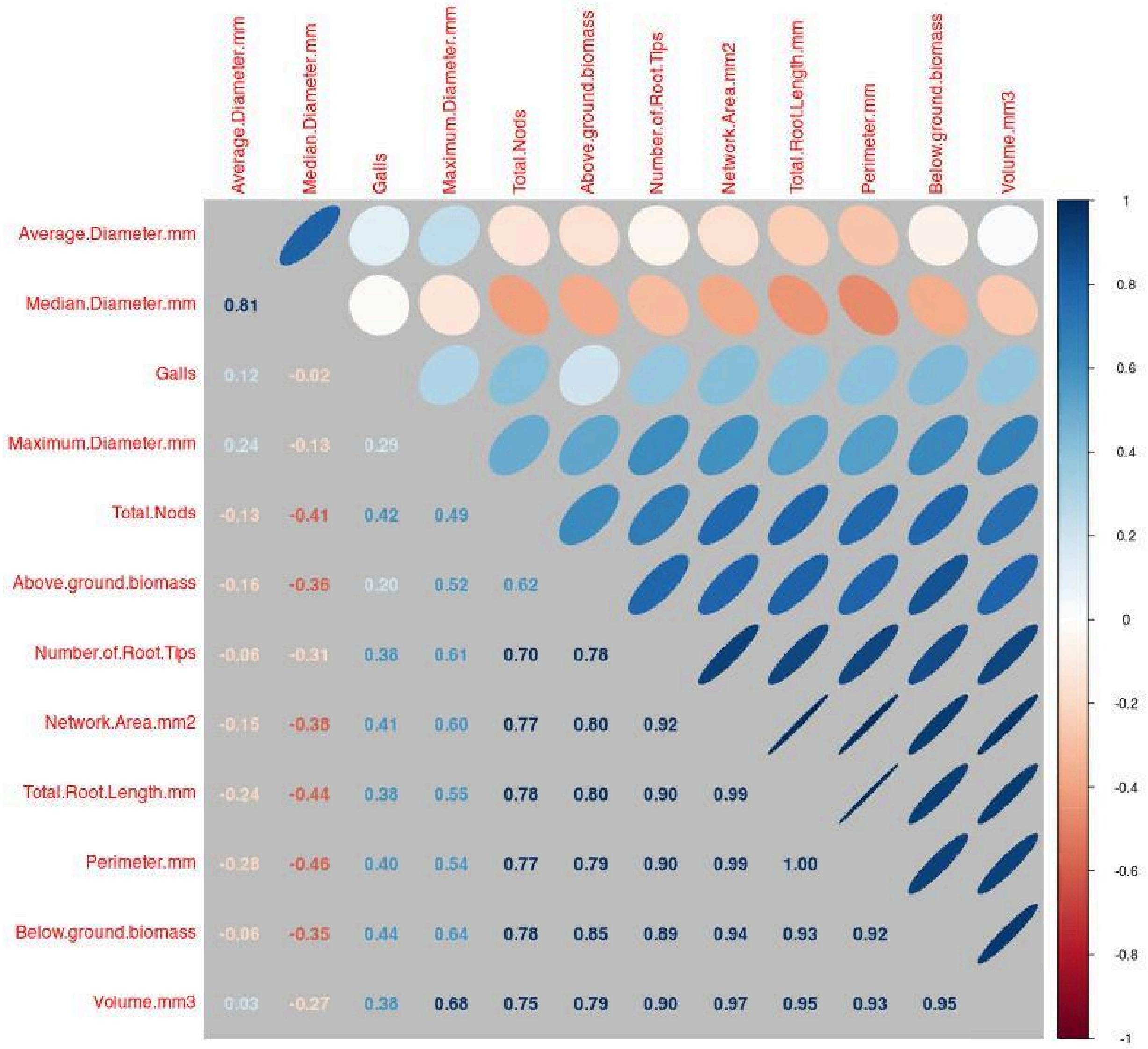

**Fig. S5**: Plot showing the correlation between root architecture traits and gall and nodule counts across all samples that were infected by nematodes.

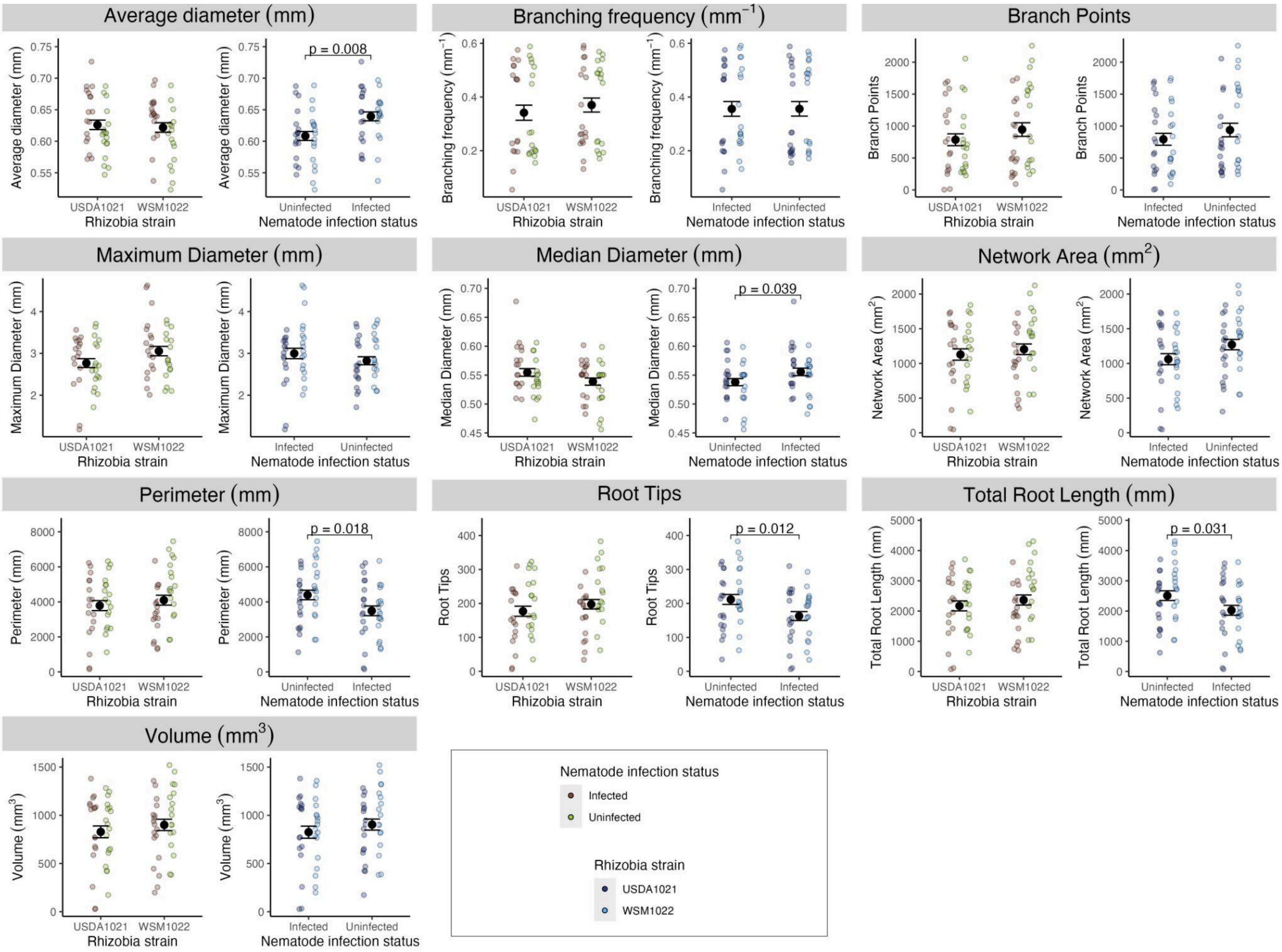

**Fig. S6**:Plots showing root architecture traits separated by nematode infection status and rhizobia strain. When statistical differences are present p values are indicated in the subplots.

**Table S1:** Gene lists of interest. See supplemental datasets for more details including log fold changes and adjusted p-values.

**a.** All genes differentially expressed in galls in response to rhizobial strain differences.

| Gene locus tag | Description (UniProt description of amino acid sequence) | Note |
|---|---|---|
| MhA1_Contig1073.frz3.gene2 | Uncharacterized protein | Nuclear localization signal sequence predicted by NLStradamus; Secreted protein predicted by SignalP 6.0 |
| MhA1_Contig1386.frz3.gene15 | Uncharacterized protein | Secreted protein predicted by SignalP 6.0 |
| MhA1_Contig253.frz3.gene16 | Cylicin-2 | Secreted protein predicted by SignalP 6.0;<br>Shows similarity to presumed parasitism gene Minc00331 in *Meloidogyne incognita* |
| MhA1_Contig338.frz3.gene3 | Amino acid kinase domain-containing protein | |
| MhA1_Contig953.frz3.gene1 | Extensin-like | Secreted protein predicted by SignalP 6.0 |

**b.** All genes differentially expressed in rhizobia strain USDA1021 in response to host parasite infection

| Gene locus tag | Description (similarity to RefSeq amino acid sequence) | Note |
|---|---|---|
| CDO29_RS04280 | metalloregulator ArsR/SmtB family transcription factor | |
| CDO29_RS13585 | Lrp/AsnC family transcriptional regulator | |
| CDO29_RS13595 | homogentisate 1,2-dioxygenase | hmgA; annotated with GO:0004411 (homogentisate 1,2-dioxygenase activity) |
| CDO29_RS14670 | phosphoserine transaminase | Annotated with GO:0004648 (O-phospho-L-serine:2-oxoglutarate aminotransferase activity) |
| CDO29_RS16270 | hypothetical protein | |
| CDO29_RS20170 | adenine deaminase | |
| CDO29_RS20175 | NCS2 family permease | |
| CDO29_RS20665 | Flp family type IVb pilin | |

| | | |
|---|---|---|
| CDO29_RS20670 | CpaF family protein | |
| CDO29_RS27030 | DHA2 family efflux MFS transporter permease subunit | Annotated with GO:0005886 (plasma membrane) |
| CDO29_RS27365 | LuxR C-terminal-related transcriptional regulator | Annotated with GO:0006355 (regulation of DNA-templated transcription) |
| CDO29_RS31060 | NADH:flavin oxidoreductase | |
| CDO29_RS31065 | dimethylsulfoniopropionate lyase | |

**c.** All genes showing similar response to rhizobial strain across all organ types. Non-putative and non-hypothetical annotations in bold.

| Gene locus tag | Description (INRAE annotation) | Note |
|---|---|---|
| MtrunA17_Chr1g0204471 | Putative protein | |
| MtrunA17_Chr2g0293681 | Putative endo-polygalacturonase | |
| MtrunA17_Chr2g0305481 | Putative protein | |
| MtrunA17_Chr2g0323741 | Putative protein | |
| MtrunA17_Chr4g0035171 | Putative glucose-1-phosphate adenylyltransferase | |
| MtrunA17_Chr4g0053411 | Putative protein kinase RLK-Pelle-LRR-III family | |
| MtrunA17_Chr5g0413411 | Putative endo-polygalacturonase | |
| MtrunA17_Chr5g0446771 | Putative Red chlorophyll catabolite reductase | |
| MtrunA17_Chr8g0364961 | **MtTPS9 (Trehalose phosphate synthase 9)** | |

**Table S2**: Select Gene Ontology terms that were enriched in various gene lists produced by differential expression analyses. See datasets for greater details regarding p and q values and other statistics.

**a.** Select enriched Gene Ontology terms within the list of genes responding to rhizobial strain identity in nematode galls.

| GO ID | Description | GO ID | Description | GO ID | Description |
|---|---|---|---|---|---|
| 0045490 | Pectin catabolic process | 0045493 | Xylan catabolic process | 0097599 | Xylanase activity |
| 0009834 | Plant-type secondary cell wall biogenesis | 0009738 | Abscisic acid-activated signaling pathway | 0010410 | Hemicellulose metabolic process |
| 0009737 | Response to abscisic acid | 0031222 | Arabinan catabolic process | 2000895 | Hemicellulose catabolic process |
| 0031221 | Arabinan meta-bolic process | 0010427 | Abscisic acid binding | 0009664 | Plant-type cell wall organization |
| 0044347 | Cell wall poly-saccharide catabolic process | 0043666 | Regulation of phosphoprotein phosphatase activity | 0071669 | Plant-type cell wall organization or biogenesis |
| 0032515 | Negative reg. of phosphoprotein phosphatase activity | 0071215 | Cellular response to abscisic acid stimulus | 0010383 | Cell wall polysaccharide metabolic process |
| 0005618 | Cell wall | | | | |

**b.** Select enriched Gene Ontology terms within the list of genes responding to nematode infection in rhizobial nodules with USDA1021.

| GO ID | Description | GO ID | Description | GO ID | Description |
|---|---|---|---|---|---|
| 0005975 | Carbohydrate metabolic process | 0008240 | Tripeptidyl-peptidase activity | 0005984 | Disaccharide metabolic process |

| | | | | | |
|---|---|---|---|---|---|
| 0006952 | Defense response | 0005576 | Extracellular region | 0022857 | Transmembrane transport activity |
| 0009311 | Oligosaccharide metabolic process | 0006112 | Energy reserve metabolic process | 0005199 | Struct. constituent of cell wall |
| 0005977 | Glycogen metabolic process | 0009737 | Response to abscisic acid | 0005506 | Iron ion binding |
| 0048046 | Apoplast | 0071944 | Cell periphery | | |

**c.** Select enriched Gene Ontology terms within the list of genes responding to nematode infection in rhizobial nodules with USDA1021 that are upregulated in uninfected plants. These terms are in addition to the terms enriched in the gene list regardless of upregulation in uninfected plants.

| GO ID | Description | GO ID | Description | GO ID | Description |
|---|---|---|---|---|---|
| 0015250 | Water channel activity | 0003333 | Amino acid transmembrane transport | 0016157 | Sucrose synthase activity |
| 0006833 | Water transport | | | | |

**d.** Select enriched Gene Ontology terms within the list of genes responding to nematode infection in rhizobial nodules with USDA1021 that are upregulated in parasitized plants. These select terms are in addition to all terms found in Table 1c.

| GO ID | Description | GO ID | Description | GO ID | Description |
|---|---|---|---|---|---|
| 0009414 | Response to water deprivation | 0043288 | Apocarotenoid metabolic process | 0006694 | Steroid biosynthetic process |
| 0006857 | Oligopeptide transport | 0005227 | Ca-activated cation channel activity | 0016125 | Sterol metabolic process |

**e.** Select enriched Gene Ontology terms within the list of genes responding to nematode infection in rhizobial nodules with WSM1022.

| GO ID | Description | GO ID | Description | GO ID | Description |
|---|---|---|---|---|---|
| 0006952 | Defense response | 0006950 | Response to stress | 0010200 | Response to chitin |
| 2001141 | Regulation of RNA biosyn-thetic process | 0080142 | Reg. of salicylic acid biosynthesis process | 0009697 | Salicylic acid biosynthetic process |
| 0006355 | Regulation of DNA-templated transcription | 0071554 | Cell wall organization or biogenesis | 0009696 | Salicylic acid metabolic process |
| 0000165 | MAPK cascade | 0007165 | Signal transduction | 0042546 | Cell wall biogenesis |

**f.** Select enriched gene ontology terms within the list of genes responding to nematode infection in rhizobial nodules in a strain-specific manner.

| GO ID | Description | GO ID | Description | GO ID | Description |
|---|---|---|---|---|---|
| 0006950 | Response to stress | 0007154 | Cell communication | 0004672 | Protein kinase activity |
| 0006952 | Defense response | 0006468 | Protein phosphorylation | 0005975 | Carbohydrate metabolic activity |
| 0031225 | Anchored component of membrane | 0044042 | Glucan metabolism process | 0006073 | Cellular glucan metabolic process |
| 0005976 | Polysaccharide metabolic process | 0010411 | Xyloglucan metabolic process | 0044264 | Cellular polysaccharide metabolic process |
| 0007165 | Signal transduction | 0051707 | Response to other organisms | 0042546 | Cell wall biogenesis |
| 0009506 | Plasmodesma | 0005618 | Cell wall | 0071555 | Cell wall organization |
| 0010411, 0046527, 0035251, | Various sugar related transferase | | | | |

| | |
|---|---|
| 0016757, 0016762, 0008194, 0016758 | activities |

**Table S3**: Table showing the constituents of the various fertilizer solutions used in this experiment.

| Solution | Molarity (mol / L) | 1 L of 0 mM N Fertilizer (mL) | 1 L of 0.625 mM N Fertilizer (mL) | 1 L of 5 mM N Fertilizer (mL) |
|---|---|---|---|---|
| $MgSO_4$ * | 0.5 | 4 | 4 | 4 |
| $CaCl_2$ * | 1 | 5 | 4.530 | 3.125 |
| Fe-EDTA | 0.02 | 2.5 | 2.5 | 2.5 |
| $MnSO_4$ | 0.006 | 0.1 | 0.1 | 0.1 |
| $CuSO_4$ | 0.006 | 0.1 | 0.1 | 0.1 |
| $ZnSO_4$ | 0.006 | 0.1 | 0.1 | 0.1 |
| $H_3BO_3$ | 0.016 | 0.1 | 0.1 | 0.1 |
| $Na_2MoO_4$ | 0.005 | 0.1 | 0.1 | 0.1 |
| $KNO_3$ | 0.625 | 0 | 0.25 | 1 |
| $Ca(NO_3)_2$ | 0.938 | 0 | 0.5 | 2 |
| $NaNO_3$ | 2.5 | 0 | 0 | 1 |
| $K_2HPO_4$ | 1.2 | 2 | 2 | 2 |
| $K_2SO_4$ * | 0.43 | 2.197 | 1.953 | 1.802 |
| NaCl | 0.2 | 1 | 1 | 1 |
| $diH_2O$ | - | 982.803 | 982.767 | 981.073 |

**Table S4**: Table showing the treatment type (nematode status and rhizobial strain) and host organ of origin for each sample.

| Sample number | Rhizobial strain | Nematode Status | Host organ |
|---|---|---|---|
| 1 | USDA1021 | Infected | Root |
| 2 | WSM1022 | Infected | Root |
| 5 | USDA1021 | Infected | Root |
| 6 | WSM1022 | Infected | Root |
| 9 | USDA1021 | Infected | Root |
| 10 | WSM1022 | Infected | Root |
| 13 | USDA1021 | Infected | Gall |
| 14 | WSM1022 | Infected | Gall |
| 17 | USDA1021 | Infected | Gall |
| 18 | WSM1022 | Infected | Gall |
| 21 | USDA1021 | Infected | Gall |
| 22 | WSM1022 | Infected | Gall |
| 25 | USDA1021 | Infected | Nodule |
| 26 | WSM1022 | Infected | Nodule |
| 29 | USDA1021 | Infected | Nodule |
| 30 | WSM1022 | Infected | Nodule |
| 33 | USDA1021 | Infected | Nodule |
| 34 | WSM1022 | Infected | Nodule |
| 37 | USDA1021 | Uninfected | Nodule |
| 38 | WSM1022 | Uninfected | Nodule |
| 41 | USDA1021 | Uninfected | Nodule |
| 42 | WSM1022 | Uninfected | Nodule |

| | | | |
|---|---|---|---|
| 45 | USDA1021 | Uninfected | Nodule |
| 46 | WSM1022 | Uninfected | Nodule |

**Table S5**: Table showing information and parameters for the DESeq2 tests performed for each model.

| Model prefix | Sample types included | Statistical effect tested for | Baseline | Genome reads were aligned to |
|---|---|---|---|---|
| med_gall_r | All galls | Effect of rhizobial strain | USDA1021 | *M. truncatula* |
| nem_gall_r | All galls | Effect of rhizobial strain | USDA1021 | *M. hapla* |
| med_nodp_r | All nodules with infected hosts | Effect of rhizobial strain | USDA1021 | *M. truncatula* |
| med_nodm_r | All nodules with uninfected hosts | Effect of rhizobial strain | USDA1021 | *M. truncatula* |
| med_root_r | All roots | Effect of rhizobial strain | USDA1021 | *M. truncatula* |
| med_nods_r21_n | All nodules with USDA1021 | Effect of nematode infection | Uninfected | *M. truncatula* |
| r21_nods_n | All nodules with USDA1021 | Effect of nematode infection | Uninfected | *E. meliloti USDA1021* |
| med_nods_r22_n | All nodules with WSM1022 | Effect of nematode infection | Uninfected | *M. truncatula* |
| r22_nods_n | All nodules with WSM1022 | Effect of nematode infection | Uninfected | *E. meliloti WSM1022* |
| med_nods_r | All nodules | Main effect of rhizobial strain | USDA1021 | *M. truncatula* |
| med_nods_i | All nodules | Rhizobial strain x nematode infection interaction effect | USDA1021, Uninfected | *M. truncatula* |
| med_gano_r | All galls and nodules with infected hosts | Main effect of rhizobial strain | USDA1021 | *M. truncatula* |
| med_gano_i | All galls and nodules with | Rhizobial strain x organ interaction effect | USDA1021, Uninfected | *M. truncatula* |

| | | | | |
|---|---|---|---|---|
| | infected hosts | | | |
| med_garo_r | All galls and roots | Main effect of rhizobial strain | USDA1021 | *M. truncatula* |
| med_garo_i | All galls and roots | Rhizobial strain x organ interaction effect | USDA1021, Uninfected | *M. truncatula* |
| med_all_r | All organs from infected hosts | Main effect of rhizobial strain | USDA1021 | *M. truncatula* |
| med_all_i | All organs from infected hosts | Rhizobial strain x organ interaction effect | USDA1021, Uninfected | *M. truncatula* |

**Table S6**: Table showing the percentage of genes in each organ and treatment type that had a coefficient of variation within the treatment type that was greater than the coefficient of variation across all samples.

| | Galls from hosts with USDA 1021 | Galls from hosts with WSM 1022 | Nodules from infected hosts with USDA 1021 | Nodules from infected hosts with WSM 1022 | Nodules from uninfected hosts with USDA 1021 | Nodules from uninfected hosts with WSM 1022 | Roots from hosts with USDA 1021 | Roots from hosts with WSM 1022 |
|---|---|---|---|---|---|---|---|---|
| USDA1021 | - | - | 4.43 | - | 11.1 | - | - | - |
| WSM1022 | - | - | - | 2.98 | - | 14.4 | - | - |
| *Medicago* | 13.6 | 12.4 | 12.7 | 13.7 | 12.8 | 15.4 | 14.5 | 19.0 |
| *Meloidogyne* | 13.1 | 16.6 | - | - | - | - | - | - |

**Table S7**: Table showing the total reads in each sample and the percentage of reads that were removed or aligned at different steps of the bioinformatic analysis.

| | s1 | s2 | s5 | s6 | s9 | s10 | s13 | s14 | s17 | s18 | s21 | s22 |
|---|---|---|---|---|---|---|---|---|---|---|---|---|
| Raw Reads | 155M | 157M | 157M | 156M | 155M | 156M | 157M | 157M | 156M | 156M | 157M | 156M |
| Total Reads after QC (sortmeRNA/ Trimmomatic) | 80M | 64M | 82M | 76M | 79M | 97M | 43M | 45M | 44M | 40M | 40M | 49M |
| Reads that aligned well (q>30; same chr) | 73M | 58M | 75M | 70M | 73M | 74M | 38M | 40M | 40M | 36M | 36M | 43M |

| | s25 | s26 | s29 | s30 | s33 | s34 | s37 | s38 | s41 | s42 | s45 | s46 |
|---|---|---|---|---|---|---|---|---|---|---|---|---|
| Raw Reads | 155M | 155M | 157M | 155M | 157M | 155M | 157M | 156M | 155M | 157M | 156M | 156M |
| Total Reads after QC (sortmeRNA/ Trimmomatic) | 83M | 74M | 78M | 84M | 84M | 79M | 82M | 90M | 76M | 85M | 84M | 84M |
| Reads that aligned well (>30 q; same chr) | 77M | 69M | 71M | 78M | 77M | 72M | 76M | 84M | 71M | 80M | 75M | 77M |

**Methods S1:** Sample preparation

Per Garcia *et al*. (2006), we mechanically scarified *M. truncatula* HM145 seeds with sandpaper, sterilized them with 10% bleach solution, stratified them in the dark at 4 °C for 2 days on sterile water agar plates, and then incubated them at room temperature for 2 days. We planted germinated seedlings in 120 mL Cone-tainers that had been filled with 1:4 mixture of perlite and sand and autoclaved twice with >24 hours between each autoclaving (Barker et al., 2006). We maintained seedlings in growth chambers at the University of Pennsylvania in a randomized complete block design of rhizobia strain (USDA1021 or WSM1022) and nematode status (N+ or N-). Each of the four treatments (USDA1021 N+, WSM1022 N+, USDA1021 N-, and WSM1022 N-) had three replicates with each replicate consisting of organ samples pooled from six individual plants. Growth chambers were kept in a 16:8 light:dark cycle, alternating between 25 °C during the light cycle and 21 °C during the dark cycle. After planting we fertilized plants on a recurring basis.

We used three different fertilizers containing 5 mM N, 0.625 mM N, or 0 mM N. Recipes for fertilizing are found in SI Table S3 and are modified from the recipes found in (Batstone et al., 2017; Moreau et al., 2008). Plants were fertilized three times a week. On the day of planting, we fertilized all plants with 5 mL of 5 mM fertilizer and 10 mL of diH2O. The three subsequent days of fertilizing we used 5 mL of 0.625 mM N fertilizer and 10 mL of diH2O. For every following day of fertilizing we used 0 mM N fertilizer and 10 mL of diH2O.

On day 14 after planting, we inoculated each *M. truncatula* plant with its corresponding rhizobia and nematode treatments. Nematode inoculation consisted of 5 mL of double deionized water solution containing 86 nematode eggs $mL^{-1}$ (~430 eggs total). Rhizobial inoculation consisted of 5 mL of 2 day old liquid culture in tryptone yeast media, diluted to OD600 of 0.1 (Journet et al., 2006). Plants not inoculated with nematodes were given 5 mL of double deionized water. Plants were fertilized on subsequent days with details of the fertilizing procedure and schedule found in SI Methods.

**Methods S2:** RNA extraction

We ground frozen organ samples within Eppendorf tubes using mini-pestles, keeping the samples frozen using liquid nitrogen the entire time. Once these samples were mostly ground, we added 1 mL of Trizol / 200mg of biomass directly to the Eppendorf tubes, and continued to grind samples until a milky semi-opaque solution formed and then vortexed the tubes until well mixed.

We heated samples at 45°C for 3 minutes minimum and then at room temperature for 5 minutes to allow the liquid to cool. We then added ~100 μL of chloroform and vortexed for 15 seconds. More than ~100 μL of chloroform was added in the previous step to get to a 1:5 chloroform:trizol ratio if needed. We closed the tubes and left them at room temperature for 2-3 minutes and then we centrifuged the tubes for 15 minutes at 4°C at 12,000g.

We transferred the resulting aqueous phase to a new tube, added 1 mL of chilled ethanol, and vortexed for 30 seconds. We then transferred the solution to a RNeasy Mini spin column and we centrifuged the column at 15,000 rpm minimum for 15 seconds, discarding what passed through the column. The column capacity required us to do the previous step 700 μL at a time, repeating as necessary. We then washed the column with 700 μL of RW1 and spun for 15 seconds at 15,000 rpm minimum, discarding what passed through the column. We then washed the column with 500 μL RPE and spun for 15 seconds at 15,000 rpm minimum, and washed with 500 μL RPE a second time but spun for 2 minutes at 15,000 rpm minimum, discarding what passed through each time.

We eluted the RNA from the column by first adding 82 μL of nuclease free H2O directly to the column and letting this sit for 5 minutes at room temperature, followed by centrifuging the column at 15,000 rpm minimum for 30 seconds while collecting what passed through in a new tube. After repeating the previous step, we added 36 μL of DNase to the elution, vortexed briefly, and then let sit the mixture sit for 25 minutes at room temperature. We then precipitated RNA by adding 3 μL of glycogen, 20 μL 3M NaOAc, and 600 μL ethanol and then vortexed the solution enough to suspend the pellet in the solution but not break up the pellet. We then put the tube in -80°C for ≥2 hours.

We centrifuged the tubes at 4°C for 90 minutes at 15,000 rpm minimum. We washed the pellet that formed at the bottom of the tube with 700 μL of 70% ethanol, and centrifuged the tube again at 4°C for 10 minutes at 15,000 rpm minimum. We poured out the ethanol solution while keeping the pellet at the bottom of the tube and opened the tubes and set them upside down to let the pellet dry for 10 minutes. We resuspended the dried pellet in 21.725 μL of nuclease-free H2O and 0.275 μL of RNase out and were put on ice for at least 30 minutes. Samples were then stored in -80°C.

**Methods S3:** Software and reference genomes

We used default settings unless otherwise stated in the Materials and Methods section. We used the following software versions and builds: samtools 1.19.2-h50ea8bc_0, Trimmomatic 0.39-hdfd78af_2, bowtie2-2.5.3-py310ha0a81b8_0, htseq-2.0.5-py310h5aa3a86_0, and sortmerna-4.3.6-h9ee0642_0.

For the reference genome of the host, we used the *Medicago truncatula* A17 ANR v5.0 genome assembly annotation release v1.9 (Pecrix et al., 2018) and used annotations from the Legume Graph-Oriented Organizer to inform our analyses (Carrï Re et al., 2020). For reference genomes for symbionts, we used the *Meloidogyne hapla* genome (Opperman et al., 2008), and the *Ensifer meliloti* strain USDA1021 and WSM1022 genomes (RefSeq ASM219744v1 and ASM1331577v1 respectively). The genomes of the two *E. meliloti* strains have a 31-kmer Jaccard distance of 0.523 or 47.3%, an average nucleotide identity of 98.27%, and a gene presence/absence Jaccard distance of 0.718 with 7,653 genes in the pangenome and 5,496 orthologous genes. To calculate similarities between rhizobial genomes, we downloaded the genomes of the two *Ensifer meliloti* strains USDA1021 (GCF_002197445.1) and WSM1022 (GCF_013315775.1) from NCBI. We computed average nucleotide identity (ANI) with fastani (ver. 1.31) (Jain et al. 2018) with default settings. We computed the Jaccard distance between genomic k-mer signatures with sourmash (ver. 4.8.4) (Titus Brown and Irber 2016) (command creating k-mer signature: sourmash sketch dna -p scaled=1000,k=31; command computing the Jaccard Index: sourmash compare --ksize 31--distance-matrix). We computed the Jaccard distance between the two genomes in terms of gene contents. This includes two steps: we annotated the two genomes using Prokka (ver. 1.14.5; options: -kingdom bacteria and --gcode 11) (Seemann 2014) and then inferred orthologous genes using Panaroo (ver. 1.3.4;options: --core_threshold 0.95 -t 10 --clean-mode strict --remove-invalid-genes) (Tonkin-Hill et al. 2020) with the mode set to strict, resulting in 5496 orthologous genes and 7653 genes in the pangenome. Scripts to calculate all these similarity metrics are also available on the github link at the beginning of this Supporting Information.

**Methods S4:** Differential expression analysis

A table outlining detailed specifications of all models used with DESeq2 can be found in supplemental materials (SI Table 4).

To test whether mutualist strain impacts gene expression at the site of parasite infection, we identified plant and nematode genes with differential expression across gall samples from plants inoculated with different rhizobia strains. To test whether nematode infection impacts gene expression in rhizobia nodules, we identified plant and rhizobia genes with differential expression across nodule samples from infected and uninfected plants. To test if nematode infection produces similar or different responses in nodules with different rhizobial strains, we identified plant genes with a main effect of nematode status and plant genes with an interaction effect of nematode status and rhizobial strain.

To determine the extent to which the effects of rhizobial strain on host gene expression are a local (i.e. unique to an individual organ) or a systemic response (i.e. similar across multiple organs), we compared the effect of rhizobia on host gene expression in galls to their effect in roots and nodules. To identify genes showing a systemic response (genes with a similar response to rhizobial strain regardless of organ type) we tested genes for a main effect of rhizobia strain. To identify genes showing a local response (genes with different responses to mutualist genotype in different organ types), we tested genes for an interaction effect of rhizobia strain and organ type.

**Results S1:** Nodule cysteine-rich protein gene expression is impacted by parasite infection

Nodule cysteine-rich (NCR) proteins are an important group of defensin-like proteins that are unique to legumes (Downie and Kondorosi, 2021). NCR proteins are highly expressed in and only in nodules (Guefrachi et al., 2014), and often play critical roles in coordinating the legume-rhizobia mutualism and nodule development and homeostasis (Burghardt, 2020; Burghardt et al., 2018; Durgo et al., 2015; Pan and Wang, 2017; Roy et al., 2020). Out of the 696 annotated NCR protein genes in the genome and annotations used in this analysis, 463 NCR protein genes (66.5%) were expressed in at least one nodule sample, and 386 (55.5%) were expressed in all nodule samples. Of all 463 expressed NCR protein genes, 74 (16%) were differentially expressed in nodules in at least one of the aforementioned models investigating nodule gene expression. Of these 74 differentially expressed NCR genes, 59 responded to nematode infection status in at least one model, 53 responded to rhizobial strain differences in at least one model with 34 responding with an interaction effect of rhizobia strain and nematode infection. Three of the differentially expressed NCR protein genes (MtrunA17_Chr3g0083081, MtrunA17_Chr3g0083091, and MtrunA17_Chr3g0083111) are present in a region of chromosome 3 that causes defective nitrogen fixation when deleted (Shen et al., 2023) and all three had a response to rhizobial strain differences or nematode infection in at least one model.
